# Moments reconstruction and local dynamic range compression of high order superresolution optical fluctuation imaging

**DOI:** 10.1101/500819

**Authors:** Xiyu Yi, Sungho Son, Ryoko Ando, Atsushi Miyawaki, Shimon Weiss

## Abstract

Super-resolution Optical Fluctuation Imaging (SOFI) offers a simple and affordable alternative to other super-resolution (SR) imaging techniques. The theoretical resolution enhancement of SOFI scales linearly with the order of cumulants, while the imaging conditions exhibits less photo-toxicity to the living samples as compared to other SR methods. High order SOFI could, therefore, be a method of choice for dynamic live cell imaging. However, due to the cusp-artifacts and dynamic range expansion of pixel intensities, this promise has not been materialized as of yet. Here we investigated and compared high order moments vs. high order cumulants SOFI reconstructions. We demonstrate that even-order moments reconstructions are intrinsically free of cusp artifacts, allowing for a subsequent deconvolution operation to be performed, hence enhancing the resolution even further. High order moments reconstructions performance was examined for various (simulated) conditions and applied to (experimental) imaging of QD labeled microtubules in fixed cells, and actin stress fiber dynamics in live cells.

## 1. Introduction

Fluorescence microscopy is widely utilized in biological studies due to its high sensitivity and specificity. These afford molecular-specific visualization of molecular structures and organelles in live cells in real-time. However, the spatial resolution of conventional fluorescence microscopy has been limited due to Abbe’s diffraction limit [1]. Advances in super-resolution (SR) imaging techniques such as stimulated emission depletion (STED) microscopy [2], photo-activated localization microscopy (PALM) [3, 4], stochastic optical reconstruction microscopy (STORM) [5], structured illumination microscopy (SIM) [6] and their derivatives allowed us to overcome the diffraction limit and achieve optical resolution down to a few tens of nanometers [3, 7-10]. Such a dramatic resolution enhancement has already yielded significant new discoveries [11-14]. A more recent addition to the SR toolbox is Super-resolution Optical Fluctuation Imaging (SOFI) [15]. SOFI has been demonstrated using different imaging platforms including wide-field microscopy (with either laser or Xenon lamp illumination) [16], total internal reflection fluorescence (TIRF) microscopy [17-21], multi-plane wide-field fluorescence microscopy [22], spinning-disk confocal microscopy [23], and light sheet microscopy [24].

SOFI relies on the stochastic fluctuations of optical signals originating from blinking emitters (see below), scatterers (as blinking Raman [25]), or absorbers [26]. Blinking fluorescence probes have been most commonly used, including fluorescent proteins (FPs) [21, 27], organic dyes [28], quantum dots [15], and carbon nanodots [19]. Other types of optical fluctuations such as ones originating from diffusion of probes [29], FRET due to diffusion [30], and stochastic speckle illumination light [31] have also been exploited for SOFI imaging.

Advantages of SOFI include compatibility with different imaging platforms and a wide variety of blinking probes, flexibility in imaging conditions [25], and a useful trade-off between spatial and temporal resolutions. SOFI has therefore the potential to democratized SR and be used in a wide variety of applications. The theoretical resolution enhancement factor for SOFI of a cumulant of order *n* is *n^1/2^* fold [15]. When combined with deconvolution or Fourier re-weighting, the enhancement factor becomes *n* fold [32]. This suggests that high-order SOFI would be beneficial for achieving high SR performance. In practice, however, two fundamental issues limit the application of high-order SOFI: (i) non-linear dynamic-range expansion of pixel intensities [15] and (ii) cusp-artifacts [33]. With regard to (i), a partial solution for the dynamic-range expansion was introduced as balanced-SOFI (bSOFI) [34]. With regard to (ii), cusp-artifacts are much harder to solve.

High order cumulants [15, 35] are constructed from correlation functions or moments of different orders. In the original introduction of SOFI [15], cumulants were chosen over moments because combinations of nonlinear cross-terms originating from multiple emitters are eliminated in the cumulants. However, as discussed in greater details in our accompanying manuscript [33] and briefly summarized here, cumulants could yield negative virtual brightnesses [33] that lead to cusp-artifacts [33]. By averaging different time blocks of cumulants, these artifacts could potentially be eliminated [36], but it requires prolonged data acquisition (with no drift) and applicable to static features only. Theoretically, another way to avoid/eliminate cusp artifacts would be to spatially manipulate emitters’ blinking behavior, so as to yield a uniform pure sign for all cumulants across the image [33]. This, however, is a very difficult task.

In this work, we examine the mathematically non-rigorous, but practical solution of moments reconstruction. We show that even-high-order moments reconstruction eliminates cusp artifacts while still providing SR enhancement. We also provide in-depth comparisons between cumulants and moments reconstruction for various simulated and experimental conditions. We also made the associated datasets [37-39] and code packages for simulation [40] and data processing [41] open to the public, as posted on the online repositories.

The outline of this manuscript is as follows: in **section 2** we briefly summarize SOFI theory and outline the relationship between correlation functions, cumulants, and moments. In **section 3**, we introduce the proposed moments reconstruction method and show that even-order moments are free of cusp artifacts. Moments reconstruction, however, introduces new artifacts due to nonlinear cross-terms. Based on the theoretical formulation, we interpret these cross-terms as contribution from *ghost emitters* in the traditional high order SOFI image [33]. We demonstrate both through theory and simulations that even-order moments yield a pure positive image, free of cusp artifacts, which is suitable for subsequent deconvolution operation. A discussion of the theoretical resolution enhancement is also discussed. In **section 4**, we introduce a new method that minimizes the ill-effects of dynamic-range expansion. We dub this method “local dynamic range compression” (*ldrc*). It locally compresses the dynamic range of pixel intensity, and its performance is not affected by cusp artifacts. This section also includes extensive simulations of various (and relevant) sample conditions that are subsequently analyzed by even-order moments reconstruction together with *ldrc*. We have compared the simulated data results with results from alternative methods, including (1) bSOFI, which utilizes the balanced cumulants to correct for the expanded dynamic range of pixel intensities of high-order SOFI cumulants, and (2) super-resolution radial fluctuation (SRRF) [42], which achieves super-resolution by calculating the degree of local gradient convergence (referred to as the radiality [42]) for each frame and characterize the corresponding temporal radial fluctuations using either temporal radiality average (TRA), or temporal radiality auto-cumulant (TRAC) of different orders. In **section 5** we present 6^th^ order moment reconstructions for experimental data together with deconvolution and *ldrc*. The data sets include quantum dot (QDs)-labeled microtubules in fixed cells and fluorescence protein-labeled β-actin in live cells. Our results are then compared to results obtained by operating the bSOFI [34] and SRRF [42] algorithms to the same data sets. A concluding discussion is given in **section 6**, summarizing our main findings: (I) even-order moments reconstruction is intrinsically free of cusp artifacts; (II) it can be independently combined with deconvolution without conflicting with the commonly used positivity constraint in image deconvolution; and (III) application of *ldrc* can correct for the expanded dynamic range of pixel intensities. These attributes allow for SR reconstruction of fast (∼seconds) morphological changes in live cells.

## 2. Review of SOFI, correlations, cumulants, and moments

We briefly repeat here the SOFI theory [15] but re-cast it in a form that affords the virtual emitter interpretation of SOFI at high orders that we proposed in our accompanying manuscript [33]. This re-casting provides insight into high order SOFI cumulants and the proposed moments reconstruction.

In the practice of SOFI, the sample is labeled with stochastically blinking emitters. This labeled sample is then imaged and consecutive frames are recorded. The data set is then SOFI processed to yield the SOFI image. Given a sample with N emitters that independently blink, the fluorescence signal captured at location 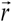 and time *t* is given by:

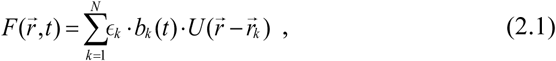

where *k* is the index of the emitter, ϵ_*k*_ is the ‘on’-state brightness of the *k*^th^ emitter, *b*_*k*_(*t*) is the stochastic time dependent blinking profile of *k*^th^ emitter where:

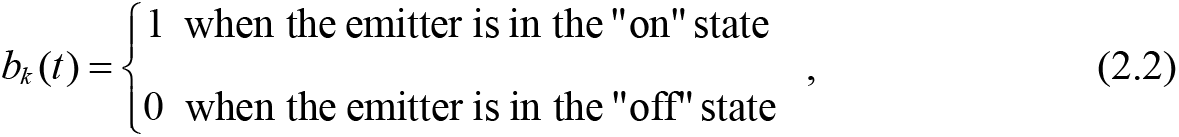

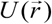 is the point-spread-function (PSF) of the imaging system, and 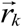 is the location of the *k*^th^ emitter. In SOFI calculations, we calculate the correlation functions along the time axis with time lags (*τ*_1_,*τ*_2_, …,*τ*_*n*_) and pixel locations 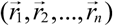:

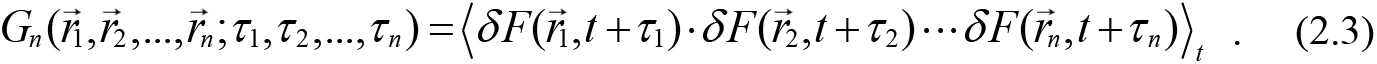

It is common to set the first time lag τ_1_ to 0, and if all the pixel locations are identical, we get auto-correlation function. Similarly, if the pixel locations are different, the correlation function is cross-correlation function. By replacing 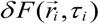 with the notation *δ F*_*i*_, equation (2.3) can be simplified:

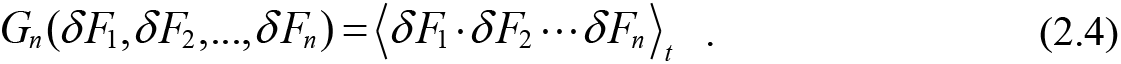

We address *G*_*n*_ (*δ F*_1_,*δ F*_2_, …,*δ F*_*n*_) as the joint correlation function for set {*δ F*_*i*_| *i* ∈[1, *n*]}, which is defined by the chosen combinations of pixels and time lags. For a given instance of time lags {*τ*_1_,*τ* _2_,…,*τ* _*n*_}, we also address *G*_*n*_ (*δ F*_1_,*δ F*_2_, …,*δ F*_*n*_) as the **joint-moment** of set {*δ F*_*i*_ | *i* ∈[1, *n*]}. The next step is to calculate the *n*^th^ order cumulant, denoted as *C*_*n*_ (*δ F*_1_,*δ F*_2_, …,*δ F*_*n*_), which we address as the **joint-cumulant** of set {*δ F*_*i*_ | *i* ∈[1, *n*]}. Note that equation (2.3) addresses cross-correlation functions, and the special case of equation (2.3) with 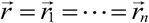 is reduced to address auto-correlation functions. Consequently, the differences between auto-correlation functions and cross-correlation functions are diminished while we form our discussion under the framework of joint-moments and joint-cumulants.

The calculation of the joint-cumulant of set {*δ F*_*i*_ | *i* ∈[1, *n*]} is illustrated in equation (2.5), using the case of 5^th^ order as an example. In the general sense, regardless of the choices of {*δ F*_*i*_ | *i* ∈[1, *n*]}, *n* fluorescence fluctuations profiles are selected from individual pixels (with or without duplicated pixel) to form the set {*δ F*_*i*_ | *i* ∈[1, *n*]} (Fig. 1 (a)), from which all the possible partitions are identified as shown in Fig. 1(b). Partitions can possess different numbers of parts where each part can possess different numbers of elements (1^st^ and 2^nd^ columns in Fig. 1(b)). For each partition, the elements of set {*δ F*_*i*_ | *i* ∈[1, *n*]} are grouped into specific parts, where each part is a subset of {*δ F*_*i*_ | *i* ∈[1, *n*]} (3^rd^ column in Fig. 1(b)). Each specific partition of set {*δ F*_*i*_ | *i* ∈[1, *n*]} contributes one term to a summation series to construct the joint-cumulant, where each term can be expressed as the product of two factors. This is shown in the 4^th^ and 5^th^ column in Fig. 1(b). The first factor *f*_1_ depends on the size of this partition (denoted as *q* in 1^st^ column in Fig. 1(b)) and is defined as: *f*_1_ = (−1)^*q*−1^ (*q* −1)! (4^th^ column in Fig. 1(b)). The second factor *f*_2_ is the product of all the joint-moments of each part within this partition, as illustrated in the 5^th^ column in Fig. 1(b): if we use *I* to represent set {*δ F*_*i*_ | *i* ∈[1, *n*]} and *I*_*p*_ (with *p*=1,2,3,…,*q*) to represent different parts that belong to this partition (as different subsets of *I*), we have *I*_1_ ∪ *I*_2_ ∪… ∪ *I*_*q*_ = *I.* The joint-moments for each part *I*_*p*_ (denote as *G*(*I*_*p*_)) are multiplied together to yield *G*(*I*_1_) ·*G*(*I*_2_) …*G*(*I*_*q*_) as the second factor (*f*_2_).

**Fig. 1.**
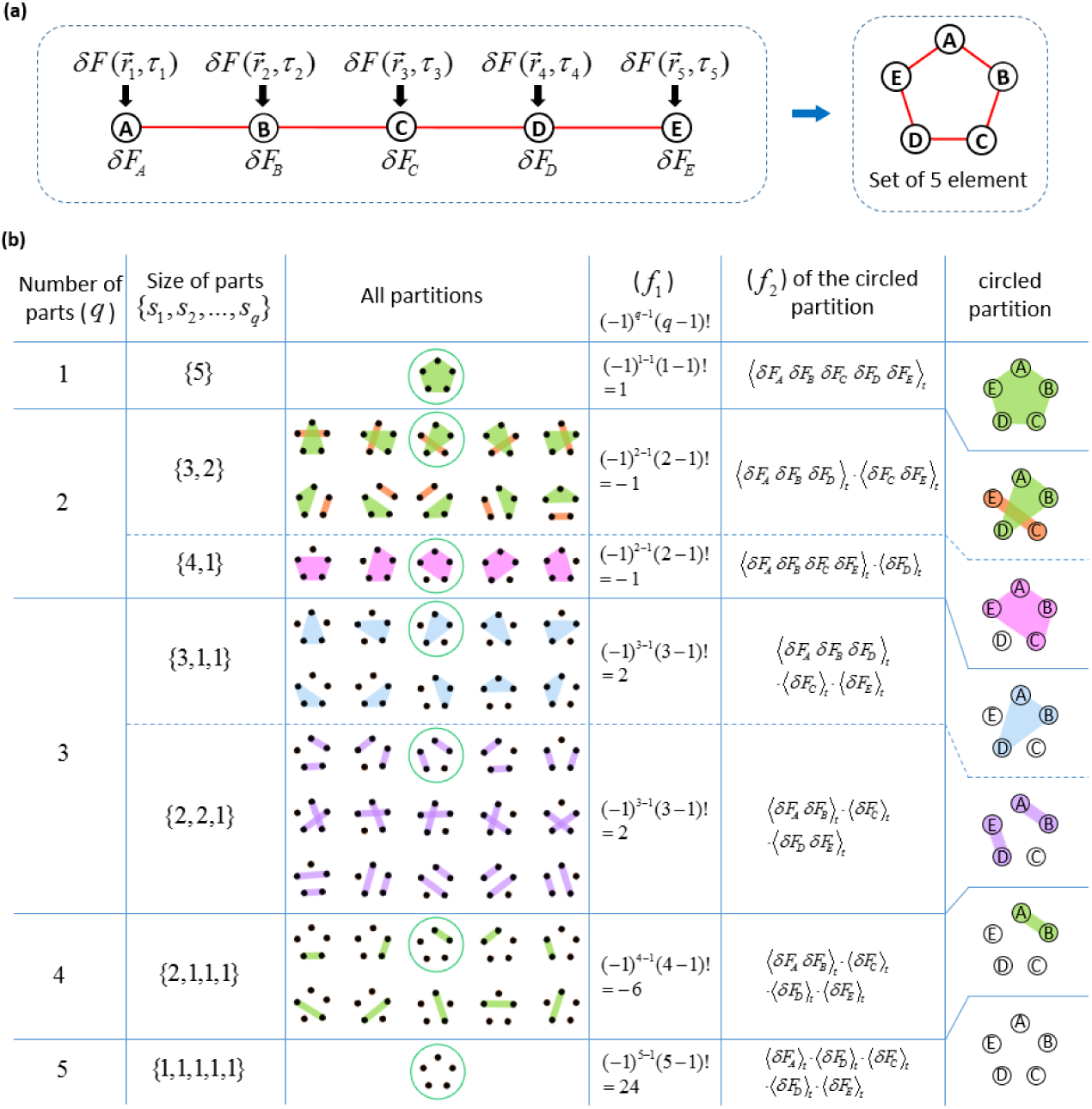
Calculation of 5^thth^ order joint-cumulants. A set of five elements is shown in (a), where the elements are the fluctuation profiles of five pixels. Repeating pixels are allowed. For example, if element A and B are repeating pixels, we have 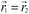 Simplified notations for the five elements are {*δF*_*A*_, *δF*_*B*_, *δF*_*C*_, *δF*_*D*_, *δF*_*E*_} respectively. (b) demonstrates all possible partitions of a set of five elements, and how each partition contributes a term to the summation series (as the product of *f*_1_ and *f*_*2*_) to yield the joint-cumulant. Note that all the partitions that contain a part of size 1 are equal to 0, because ⟨*δ F*(*t*)⟩_*t*_ = 0. The graphical demonstration of partitions is inspired by the work by Tilman Piesk [43].

In conclusion, given a set of intensity trajectories from a group of pixels (set *I*) (either with or without duplicated pixels), the joint-cumulant of *I* is constructed as a function of the joint-moments of all parts over all possible partitions of set *I*, based on the following formula [44]:

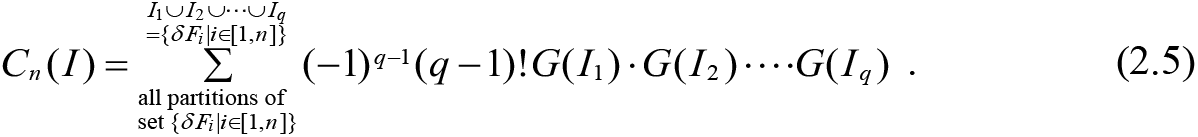

Note here that in equation (2.5) above, the joint-moments *G*(*I*_*p*_) are essentially the lower order correlation functions discussed in the original SOFI paper [15]. If a partition contains a part that has only one element, we have the corresponding *G*(*I*_*p*_) as ⟨*δ F* (*t*) ⟩_*t*_ = 0. As a result, the corresponding *f*_2_ factor will be 0, and this partition will not contribute to the joint-cumulant. The calculation of *C*_5_(*I*) is shown in Fig. S1 [45] as an example.

By substituting equation (2.1) - (2.4) into equation (2.5), we find that the *n*^th^ order joint-cumulant of set {*δ F*_*i*_ | *i* ∈[1, *n*]} can be expressed as follows:

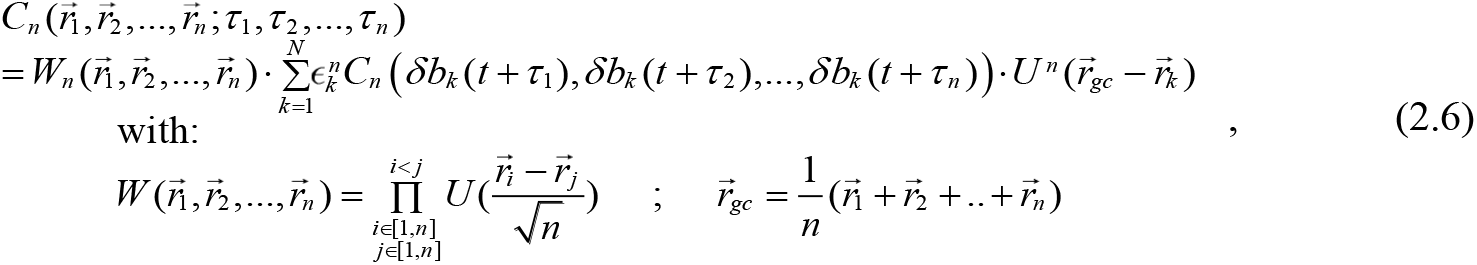

where 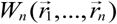 is the distance factor [15]. The PSF can be approximated by a Gaussian:

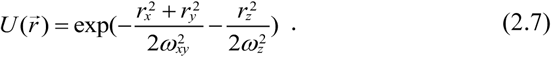

Detailed derivation of equation (2.6) can be found in Appendix 1 [45]. Once the distance factor is solved and divided from both sides of equation (2.6), the cumulant value at location 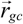 is obtained.

The SOFI pixel location vector is equivalent to the vector average of the selected pixels’ locations (in case of pixel repetitions, repeat the corresponding location vectors as well). The choice of pixel combination imposes a trade-off between noise contribution and the attenuation imposed by the distance factor 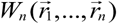 (defined in (2.6)). On one hand, noise could potentially contribute to the resultant cumulant value if there is pixel repetition in the selection. On the other hand, when the selected pixels are distributed too far away from each other, the distance factor becomes small and attenuates the correlation signal. Existing approaches have been focused on avoiding the noise contribution from duplicated pixels [46], but here we explore and present the opposite of this trade off, where we want to diminish the effect of the distance factor at the cost of potential noise contribution. A detailed explanation for our choice of pixel combinations for high order SOFI is given in Appendix 2 and Fig. S2 [45].

Under the framework of virtual emitter interpretation [33], the physical meaning of the joint-cumulant calculated for a set of pixels (either with or without pixel repetition) is taken to mean as the image formed by virtual emitters at the locations of the original emitters, but having virtual brightnesses. These virtual brightnesses are the products of ϵ^*n*^ (meaning the *n*^th^ power of the original ‘on-state’ brightness of the emitter) and *w*_*n*_(δ*b*_*k*_(*t*)) (meaning the *n*^th^ order cumulant of the blinking profile of the corresponding emitter with the time lags defined for the overall joint-cumulant function). Because the blinking statistics of emitters across the image are not necessarily spatially uniform, the ‘on-time ratio’, defined as the percentage of time the emitter spent at ‘on’ state, can vary, causing cumulant values to have different signs at different parts of the image (Fig. 2). Since images are usually presented with positive pixel values, the absolute value operator could yield an image with cusp-artifacts, degrading the image quality of high-order SOFI cumulants [33].

**Fig. 2.**
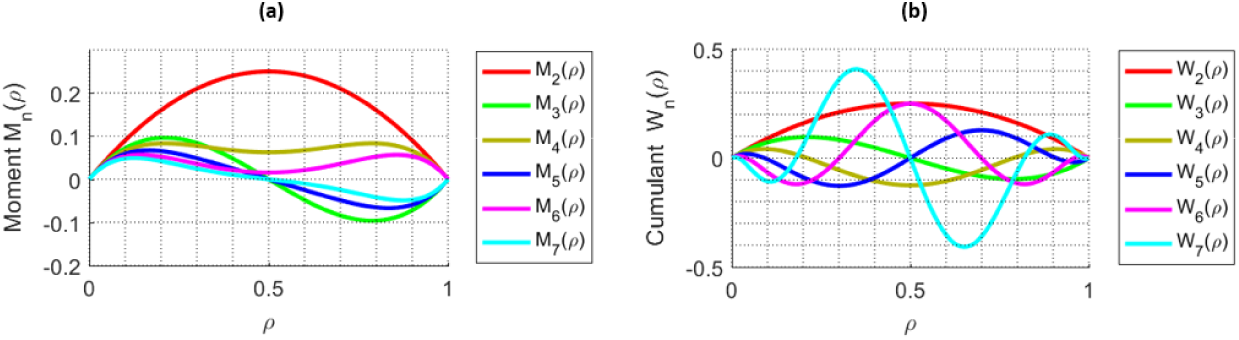
Moments and cumulants as a function of the ‘on time ratio’ ρ. (a) shows different moments as a function of ρ and denoted as M_n_(ρ), and (b) shows different cumulants as a function of ρ and denoted as W_n_(ρ). In both notations, *n* represents the order.

## 3. High-order moments reconstruction – theory and Interpretation

Inspired by the interchangeable relation between cumulant and moment [35], we investigated the statistical behavior of high-order moments of emitter blinking trajectories expressed as a function of the ‘on time ratio’ ρ in a similar way to cumulant analysis [33]. Considering only the blinking profile (with unit brightness) as shown in equation (2.2), the ‘on’ state signal is 1, the ‘off’ state signal is 0, the time average of the blinking trajectory is ρ, therefore, after subtraction of the time average ρ, the blinking trajectory exhibits (1-ρ) and (-ρ) for the ‘on’ and ‘off’ states respectively. We call this signal after subtraction of the time average as the ‘center-shifted’ signals. Additionally, the percentage of ‘on’ and ‘off’ state in the overall trajectory is ρ and (1-ρ) respectively, providing the weighting factors for both states when calculating the moments. The *n*^th^ order moment can be readily calculated as the weighted summation of the *n*^th^ power of the ‘center-shifted’ signal for both states:

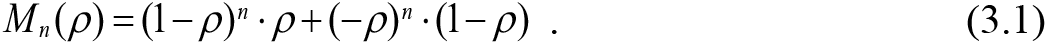

Figure 2 shows moments of different orders as a function of ρ (Fig. 2(a)) in comparison to cumulants of different orders as a function of ρ (Fig. 2(b)). While cumulants exhibit oscillation between positive and negative values, even-order moments have pure positive values (and odd-order moments are bi-modal with a single node).

In practice, blinking behavior of fluorophores are often not well controlled or adequately sampled, therefore, can be composed of mixtures of positive and negative virtual brightnesses [33], leading to cusp artifacts [33]. Since even-order moments are always positive and could therefore eliminate cusp artifacts, we decided to examine their ability and fidelity in reconstructing SR images of high order. As explained in the introduction, such a reconstruction is mathematically non-rigorous due to nonlinear cross-terms containing mixed signals from multiple emitters. Our examination could, however, evaluate the benefits of eliminating cusp artifacts vs. the drawbacks of introducing additional virtual emitters (originated from cross-terms). Moreover, since even-order moments reconstruction contains pure sign (purely positive), and the absolute image is free of cusp artifacts, subsequent deconvolution operation (that often carries positivity constrain) could further enhance the resolution.

In order to better understand the physical meaning of moments, we look at the form of moments derived from cumulants [35]:

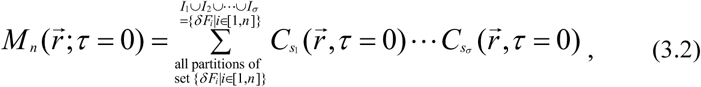

where {*I* _*p*_ | *p* ∈[1,*σ*]} is one partition of set {*δ F*_*i*_ | *i* ∈[1, *n*]}, σ is the size of this partition (i.e. the total number of parts within this partition), *S*_*p*_ is the size of *I*_*p*_, and 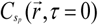 is the 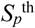 order cumulant of fluorescence fluctuation at location 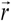 with all the time lags equal to 0. Note here that we use σ to represent the partition size instead of using *q* to distinguish moments reconstruction from cumulants reconstruction. The reconstruction algorithm is shown in the flow diagram of Fig. S3 [45]. With the goal of achieving *n*^th^ order moments reconstruction, we can interpolate all calculated cumulants (2^nd^ order to *n*^th^ order) onto a unified high resolution spatial grid that supports all orders. This re-mapping provides a full set of cumulants for each pixel if we need interpolation. Next, different orders of cumulants are combined (as shown in equation (3.2)) to reconstruct the moments at each pixel, thus achieving the *n*^th^ order moment reconstruction. A similar re-mapping could be achieved using fSOFI [47] with interpolation performed on each individual frame of the acquired before correlation calculations to directly compute moments. When all the time lags used in the correlation calculation are 0, the computational cost for the moments computation at each interpolated pixel is greatly reduced. The analytical expression for the reconstructed moments can then be expressed as:

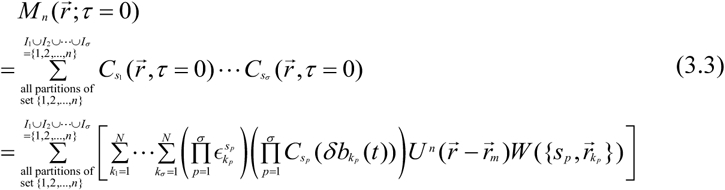

where *W* is the ‘emitter distance factor’, whose analytical form is the same with that of the distance factor [15], and is dependent on the mutual distances between different pixels as shown in (2.6). Detailed derivations of equation (3.3) is given in Appendix 3 [45]. We also define 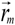 as:

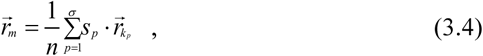

to be the mass center of the mass points (indexed with *p* as shown in (3.4)) at locations 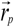 with mass values *S*_*p*_. We can re-index the summation series of equation (3.3) into the summation over all possible mass centers. Consequently, the moments reconstruction is formed as the convolution between a virtual PSF 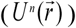 and a virtual ground truth location map constitutes of all the mass centers. The virtual PSF is the original PSF raised to the power of *n* that maintains the theoretical resolution enhancement, and the virtual ground truth location map is described by superposition of virtual emitters with locations described by (3.4). To gain more intuitive insight, the summation series in equation (3.3) can be divided into two parts. The first part is the case when all the emitter vectors in (3.4) are the same, they describe the virtual emitter that is located at the original real emitter location. As shown below:

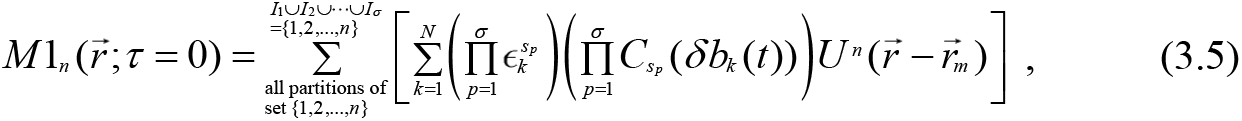

*M*1_*n*_ is the part with identical location vectors representing real emitters at locations 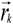. The equation can be simplified into the following form (as shown in Appendix 8 [45]):

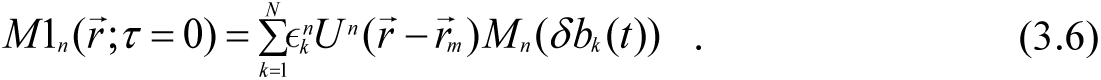

From equation (3.6) we deduce that this portion of the signal (*M*1_*n*_) is equivalent to an image formed by virtual emitters that are located at the same locations as the original emitters with brightnesses 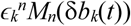 (for the *k*^th^ virtual emitter). These brightnesses differ from the ones derived for cumulants [33]: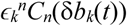. For the *k*^th^ virtual emitter, its virtual brightness is the product between the *n*^th^ power of its on-state brightness 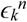 multiplied by the *n*^th^ order moment (instead of cumulant) of its blinking fluctuation δ*b*_*k*_(*t*). Because 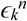 is always positive, even order moments are always positive, therefore the virtual brightness for this portion of the moments signal are always positive.

The second, non-physical part of the summation series in equation (3.3) is the case where the partitions contains non-identical emitter location vectors. The corresponding virtual emitters are located at locations where there are no real emitters (unless by coincidence the mass center overlaps with the location of a real emitter with rare chances). This part represents additional virtual (artificial) emitters at locations vectors that are not identical. It originates from cross-terms of signals coming from non-identical emitters. They take the form of virtual emitters at new locations (different from locations of original real emitters; dubbed here as ‘ghost’-emitters). The brightnesses of these ‘ghost’-emitters are attenuated by the emitter distance factor, ranging from 0 to 1 as represented in the same analytical form of the original distance factor [15].

## 4. High-order moments reconstruction of simulated data

To take a close-look of the resolution enhancement and assess the contribution of ‘ghost’-emitters, we simulated 3 near-by Poisson-blinking fluorophores and calculated the moments of the simulated movie (Fig. 3). The parameters used to generate the blinking trajectories are tabulated in Fig. 3(a), and the positions of the 3 emitters are shown in Fig. 3(b). In Fig. 3(c), the 6^th^ order moments are compared with the theoretical prediction, which is calculated from equation (3.3) from the ground truth parameters. The resolution enhancement also confirmed in Fig. 3(d) with increasing order of moments and decreasing size of the PSFs of the three emitters. We note that the prediction is affected by the time-binning introduced by the camera’s integration time for each frame. A correction for the binning effect could be introduced to the theoretical framework as was done by Kendall et al [35] (but this is beyond the scope of the work presented here).

**Fig. 3.**
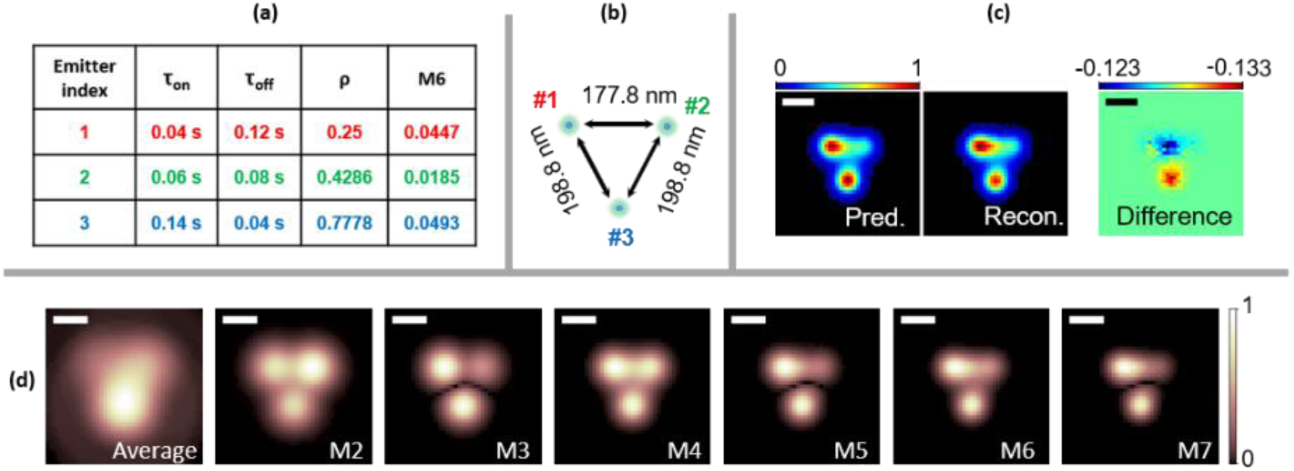
Moments reconstruction of simulated data for 3 near-by blinking fluorophores. (a) shows the photophysical parameters used in the blinking simulation of the three emitters. (b) shows the ground truth location of the three emitters. Other parameters used for the simulations: emission wavelength of 520nm, numerical aperture of NA=1.4, frame integration time of 2 ms. The pixel size was set to be small (17.78 nm) to avoid artifacts due to binning. (c) shows the comparison between the prediction (Pred.) and reconstruction (Recon.) of the 6^th^ order moment. (d) shows the average image (Ave.) and moments of the simulated movie (M2 to M7). Scale bars: 160 nm.

Besides, equation (3.3) indicates that the emitter distance factor *W(s*_*p*_, *r*_*k*_) attenuates the virtual brightnesses of the ghost emitters, and their contribution to the image is to ‘fill-in’ the space in between of the real emitters #1, #2 and #3. Such insight is confirmed in Fig. S6 and Fig. S5 [45], where the distances of the three emitters are progressively increased, and the ghost emitters’ intensities are more attenuated with the increase of such increased distances (Fig. S6 [45]), indicating that the ‘ghosting’ effect is a near-range effect, and Fig. S6 indicates that the critical distance is between 160 nm to 177.8 nm. We further simulated two lines that are placed in parallel at a distance of 160 nm. Results are shown in in Fig. S7 [45], with different total number of frames, labeling densities and pixel sizes. Comparisons are performed between the average, 6^th^ order cumulants with and without taking the absolute value, and 6^th^ order moments with and without *ldrc*. ‘Ghost’ emitters in between the two lines are attenuated because of the relatively large distance. ‘Ghost’ emitters along the same line are not much less attenuated, but contribute to the overall ‘smoothing’ of the filamentous feature. Nonetheless, despite this smoothing-out, SR enhancement is still maintained, as shown in Fig. S7 [45]. The two lines are resolved in the 6^th^ order moments reconstruction, even with larger pixel sizes and less frames. Fig. S7 [45] shows that using less total number of frames and larger pixel sizes can negatively impact both cumulants and moments, but moments reconstruction is more robust as compared to cumulant reconstruction.

Considering the existence of ghost emitters, the limit of the resolution enhancement of *n*^th^ order moment reconstruction (where *n* is an even number) is the resolution enhancement acquired in the 2^nd^ order moment (equivalent to the 2^nd^ order cumulant in our case): 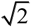. However, even order moments are strictly free of cusp-artifacts (see Fig. 2(a)), deconvolution algorithms could be readily applied to further enhance the resolution by up to an addition of *n*^1/2^ fold [32], resulting in a theoretical 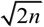 fold resolution enhancement. This resolution enhancement factor is higher than that for pure cumulants without deconvolution (*n*^1/2^), but lower than that for cumulants with deconvolution (*n*), but cusp-artifacts greatly corrupt the high-order cumulants, rendering such deconvolution impractical. A similar argument holds for bSOFI reconstructions which assumes perfect deconvolution. In summary, the artifacts introduced by ‘ghost’ emitters in moments are manageable. Even-order moments indeed exhibit ‘ghosting’ artifacts, but they are limited due to the brightness attenuation. Importantly, even-order moments are free of cusp artifact because virtual and real brightnesses are positive, allowing for a subsequent deconvolution step that improves the total resolution enhancement of up to a factor of 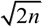.

We further increased the complexity of the simulations for various sample conditions and assessed the performance of the 6^th^ order moments reconstructions in comparison to the performance of bSOFI and SRRF reconstructions for the same data sets. bSOFI is designed to solve the dynamic range expansion of the high order SOFI cumulants, but assumes perfect deconvolution, rendering the method vulnerable to the cusp artifacts. And SRRF calculates radial fluctuations, exhibiting as an alternative SR method that utilizes fluctuations information of the acquired image. The moment reconstruction was combined with a cusp-independent dynamic range compression method termed ‘local dynamic range compression (*ldrc*)’. In ldrc, the dynamic range of the pixel intensities of high order SOFI image is compressed in a local manner with a lower order SOFI image serving as a reference image. Specifically, a small window (typical width is 35 to 75 pixels) is defined, and the pixel intensities within the window are rescaled to the range of the same windowed area in the reference image. The window is moved across the field of view while each windowed area is rescaled independently. The output image with compressed dynamic range is the average of all the rescaled windows. The reference image is usually the 2^nd^ order SOFI image that has excellent background removal and moderate dynamic range expansion. A more detailed discussion about *ldrc* is provided in Appendix 4 [45]. All reconstructions were compared to the ground truth image. As shown in Fig. 4, bSOFI reconstructions exhibits discontinuities in the simulated filaments while SRRF artificially narrows them down. moments reconstructions yield a more faithful representation of the simulated data as compared to the ground truth.

**Fig. 4.**
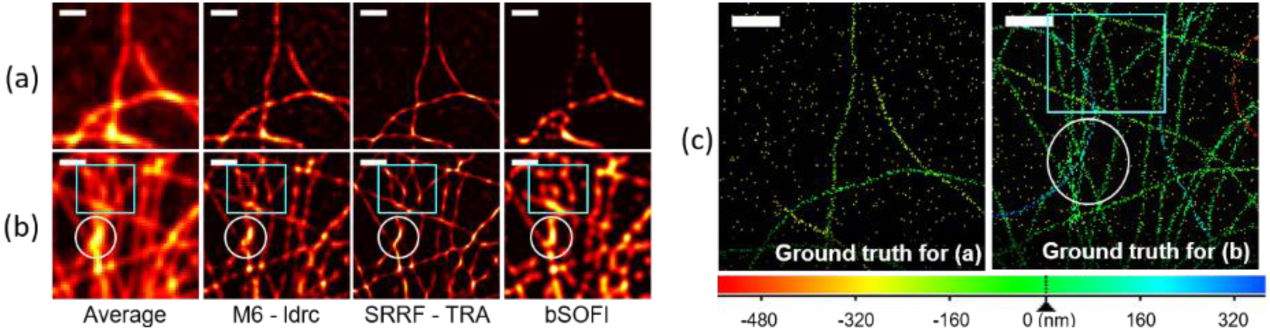
Comparison of high-order moments reconstruction with high-order bSOFI and SRRF reconstructions on simulated filaments. A simulated dataset consisting of filaments in a 3D space was generated with: 50 emitters per 1 um labeling density along the line, 10 nm cross-section thickness with a Gaussian profile, 520 nm emission wavelength, 1.4NA and 90x magnification and a grid of 125×125 pixels with a pixel size of 1.6 × 1.6 um^2^. The Gibson Lanni’s PSF model was used in the simulations. Small field of views are cropped with different feature densities for comparison. (a) Sparse filaments. All methods yield satisfactory results. While M6-ldrc exhibits some grids artifacts, SRRF emphasizes thin features with oscillatory intensities and bSOFI exhibits granular and discontinuous features. (b) Dense filaments. Compared to the ground truth image, M6-ldrc exhibits the most faithful representation, while SRRF-TRA omits filaments (circled area for example). bSOFI exhibits discontinues filaments and features at locations that have no ground-truth signal (boxed area for example). (c) and (d) shows the ground truth for (a) and (b) as labeled in the image respectively. Scale bars: 640 nm.

Further reconstructions results for a variety of simulated challenging image conditions are summarized in as supplementary figures in the appendix [45], including for different labeling density (Fig. S8 [45]), increased filaments thickness (or equivalently labeling uncertainty) (Fig. S9 [45]), increased nonspecific binding emitter density (Fig. S10 [45]), and various signal levels (Fig. S11 [45]).

Details of the simulations are given in Appendix 5 [45]. We further tested the 3D sectioning capability on an additional set of simulations where acquisitions of the same simulated sample at 100 different focal planes were generated [37] and processed independently and subsequently combined for 3D reconstruction. *ldrc* together with moments reconstruction have yielded better sectioning performance than SRRF when compared to the ground truth of the simulation (Fig. 5, Visualization 1, and Visualization 2).

**Fig. 5.**
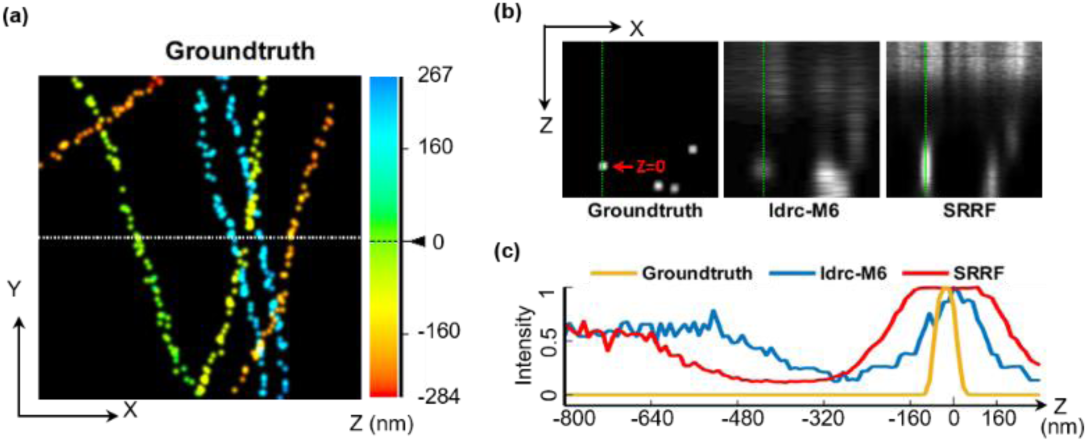
Comparison of high-order moments reconstruction with high-order SRRF reconstruction for 3D sectioning performance. 3D sectioning results of ldrc-M6 and SRRF on simulated data are shown for a small field-of-view (2.15 μm × 2.15 μm). The full field-of-view results during a continuous scan of of the focal plane is provided in SI Movie 1. (a) shows the ground truth image of the simulated filaments projected onto x-y plane. Emitters are represented by 3D delta functions convolved with a 3D Gaussian with FWHM = 86.27 nm for the purpose of display. The color scale represents the *z* coordinate of the emitters. (b) x-z scan corresponding to the dashed line in (a), where 4 filaments are penetrating through the plane (a fifth filament (yellow) is missing at this plane because the sparse, stochastic labeling algorithm did not place an emitter at the corresponding (*x, y, z*) coordinate. (c) A z-direction cross section of the first (green) filament for ground-truth and ldrc-M6 and SRRF reconstructions.

## 5. High-order moments reconstruction of experimental data

High-order moments reconstruction (6^th^ order) in combination with *ldrc* and deconvolution were applied to experimental data of quantum dots-labeled α-tubulin filaments in fixed Hela cells. The results are compared to bSOFI and SRRF results (Fig. 6). As shown already in the previous section, SRRF exhibit the highest visual resolution enhancement, but at the expense of introduction of distortions, while ldrc-M6 exhibits more faithful results (to the average image).

**Fig. 6.**
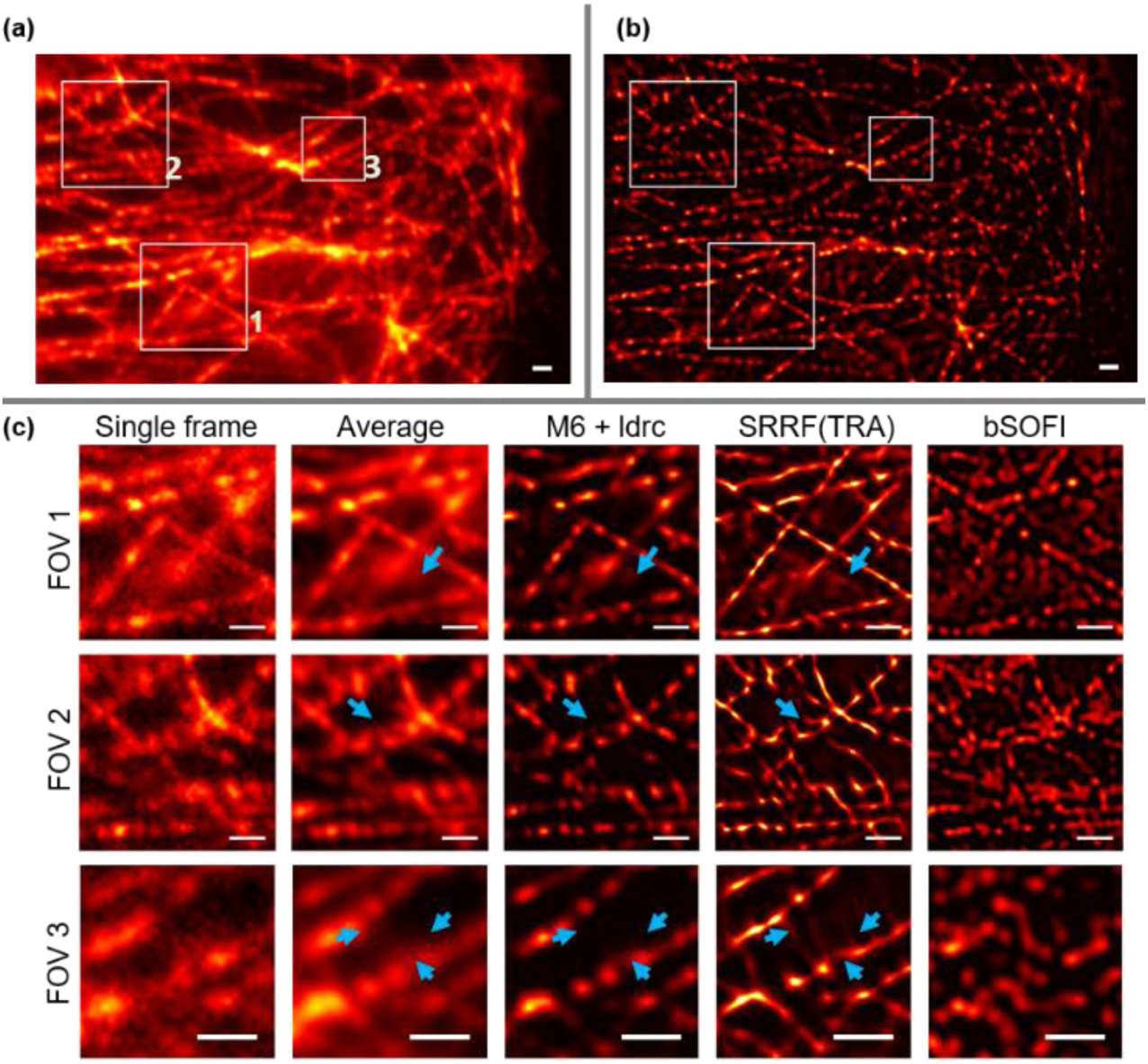
High-order moments reconstruction of experimental data (fixed cells). α-tubulin filaments in fixed Hela cells were labeled with QD800. 1000 frames were acquired with 30 ms integration per frame and processed. (a) shows the average image (b) shows the ldrc-M6 results from the full field-of-view. Three zoom-in panels in (a) are shown in panel (c) as FOV1, FOV2 and FOV3 respectively, for single frame, average image, and results from ldrc-M6, SRRF and bSOFI respectively. The displayed SRRF result is optimized among options of TRA and TRAC of different orders. bSOFI exhibit discontinuities, SRRF provides higher resolution details but with distortions (blue arrows) and extra features which could be perceived as extra filaments with perpendicular branching angle to the robustly visualized microtubules. However, the microtubule branching angles are most commonly distributed within a range of 20° to 60 ° with average of 40° [48], the perpendicular branching displayed in the SRRF could be an artifact. The ldrc-M6 image is similar to the average image but with less background and improved resolution. Scale bars: 800 nm.

The ldrc-M6 results for live cell imaging [38] are shown in Fig. 7. Fluorescence labeling was performed by fusing β-Actin protein sequence to either Skylan-S [21] or Dronpa-C12 (Fig. S16 [45] and Appendix 6 [45], R. A. et al., manuscript in preparation) with a (GGGGS)×3 linker. The bSOFI algorithm does not perform particularly well for this frame rate (33 Hz). SRRF, on the other hand, exhibits excellent performance in terms of resolution enhancement and highlighting and preserving small features (green arrows), but at the cost of introducing extra features that could be artifacts (blue arrows). In addition, M6 results afford deconvolution post processing (DeconvSK [49]), while deconvolution performed on SRRF results highlights the artifacts. The reproducibility of the reconstruction algorithms and their comparisons could be assed from reconstructions of additional experimental data sets (Fig. S12, S13, S14, S15 [45] and and Visualization 3, Visualization 4, Visualization 5, Visualization 6, Visualization 7, Visualization 8). Details of the experiments can be found in Appendix 6 [45], and details of data processing can be found in Appendix 7 [45]. Both SRRF and Moments reconstruction (M6+ldrc+deconvolution) outperform bSOFI and SOFI cumulants, especially when applied to fast live cell imaging data.

**Fig. 7.**
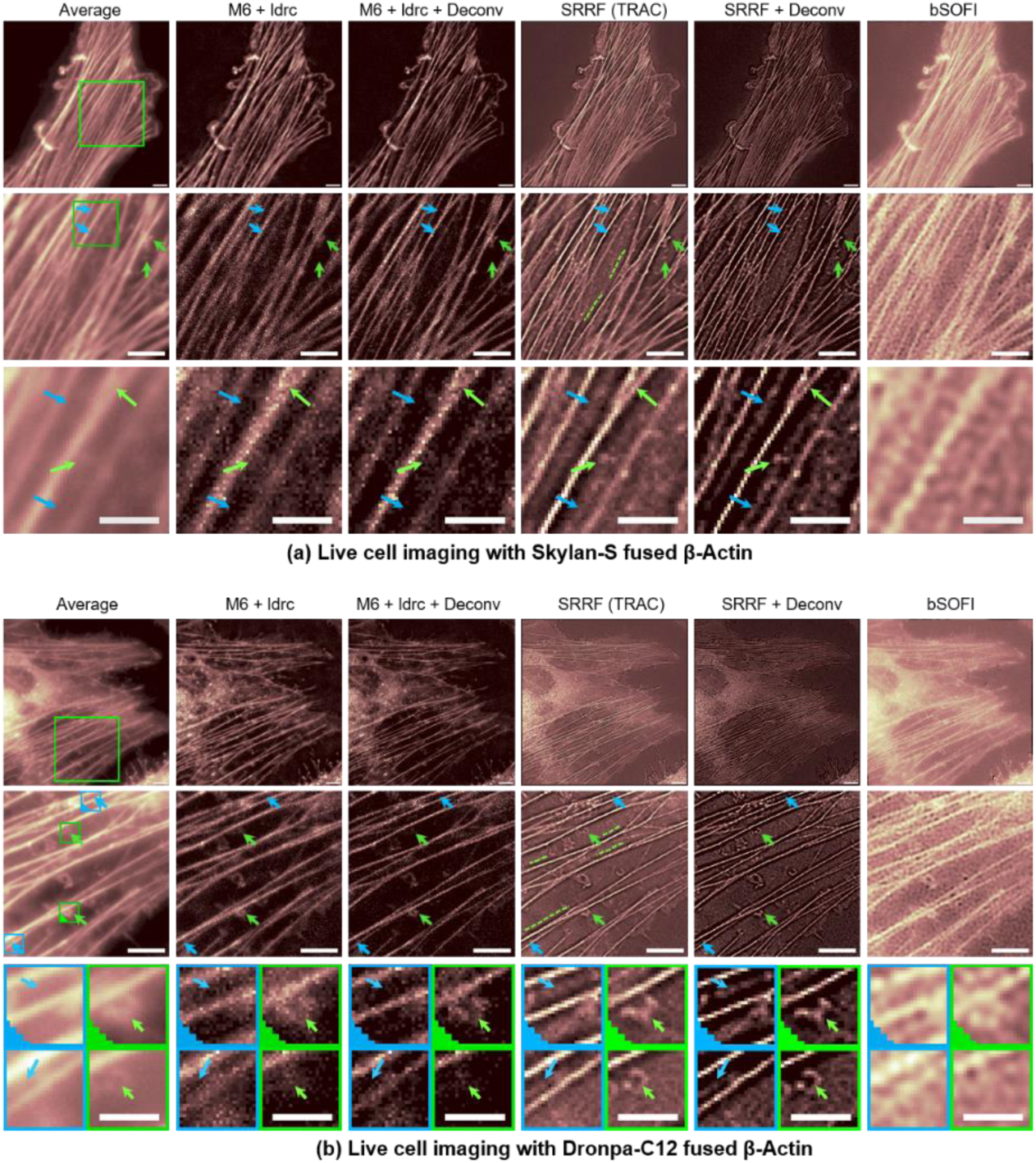
High-order moments reconstruction of experimental data (live cells). Hela cells were transfected with plasmid encoding (a) Skylan-S fused to β-Actin and (b) Dronpa-C12 fused to β-Actin. Live cells were imaged with 30 ms frame integration. 200 frames of the movie were processed per block. For each panel, the top row shows the full field-of-view, the middle row shows a zoom-in region (green box in the Average image), and the bottom row shows the further zoom-in region (green and blue boxes in the middle row image with or without a triangle marker at the left bottom corner). Each column shows results for the reconstruction method labeled at the top. We can see that while SRRF exhibit excellent performance on highlighting small features (green arrows), but at the cost of introducing shadow artifacts (along and under the green dashed lines) along the sides of bright filaments, and extra feature that could be artifacts (blue arrows). The displayed SRRF result is optimized among options of TRA and TRAC of different orders. Scale bars: 8 μm.

## 6. Conclusions

As explained in our accompanying work [33], cusp artifacts greatly affect the quality of high-order SOFI (cumulant) reconstruction. In this paper we reexamined the mathematically non-rigorous moments reconstruction and compared its results with SRRF and bSOFI reconstructions. Moments reconstructions (combined with *ldrc* and deconvolution) of simulated and experimental data sets exhibited satisfactory results with resolution enhancement and minimal distortions. Although they inherently introduce additional, spurious signals from the ghost emitters, in practice, the reconstructions are faithful to the ground-truth of simulated data and average image of experimental data. Moments reconstruction and SRRF both outperform bSOFI due to the latter’s heavy reliance on deconvolution. In contrast to bSOFI, Moments reconstruction allows for the subsequent application of deconvolution to the reconstruction, independent of the dynamic range compression process. The theoretical resolution enhancement factor for even order moments is at least 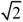, and once combined with subsequent deconvolution algorithm, such factor can be improved to 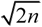. Lastly, we have demonstrated a super-resolved M6-reconstructed live cell movie with a temporal resolution of 6 seconds per frame (requiring only 200 frames of the original movie for each frame of the reconstructed movie) using a conventional wide field fluorescence microscope.

## Supporting information

Visualization 1

Visualization 2

Visualization 3

Visualization 4

Visualization 5

Visualization 6

Visualization 7

Visualization 8

## Acknowledgement

S. W. and X. Y. designed the research, X. Y. and S. S. performed the experiments, X. Y. developed simulations, the analysis methods and the associated codes, R. A. and A. M. designed and developed the Dronpa-C12 protein. S. W. and X. Y. wrote the manuscript. These authors declare no conflict of interests. We would like to thank Prof. Pingyong Xu for providing the Skylan-S plasmid. We also would like to thank Ms. Yingyi Lin and Mr. Xi Lin for their help with live-cell experiments as undergraduate student researchers.

