## Appendix for "Moments reconstruction and local dynamic range compression of high order superresolution optical fluctuation imaging"

### Appendix 1

This note derives equation (2.6) by substituting equations (2.1) to (2.4) into (2.5) (see main text). We start with equation (2.5):

$$C_n(I) = \sum_{\substack{I_1 \cup I_2 \cup \cdots \cup I_q = \{\delta F_i | i \in [1,n]\} \\ \text{all partitions of set } \{\delta F_i | i \in [1,n]\}}} (-1)^{q-1}(q-1)! G(I_1)\cdot G(I_2)\cdots G(I_q) \quad , \qquad \text{(SI-1.1)}$$

where $G_n(I)$ is the n<sup>th</sup> order cumulant of set $I$ that contains n elements $\{\delta F_i(\vec{r}_i, t+\tau_i) \,|\, i \in [1,m]\}$, and the summation over all possible partitions of set $I$ where each partition is represented as a set of subsets of $I$ such that $I_1 \cup I_2 \cup \cdots \cup I_q = I$, and $q$ is the total number of parts of this specific partition. Consider $I_p = \{\delta F_i(\vec{r}_i, t+\tau_i) \,|\, i \in [1,m]\}$ as one subset of a partition of $I$, where $m$ represents the size of this subset. $G(I_\mathrm{p})$ would then represents the cross correlation of all the elements that belongs to $I_p$. Consider the analytical form of $\delta F_i(\vec{r}_i, t+\tau_i)$:

$$\delta F(\vec{\gamma}_i, t+\tau_i) = \sum_{k=1}^{N} \epsilon_k \cdot \delta b_k(t+\tau_i)\cdot U(\vec{\gamma}_i - \vec{r}_k) \quad , \qquad \text{(SI-1.2)}$$

where $\vec{\gamma}_i$ is the location of the $i^{\text{th}}$ pixel having a fluorescence fluctuation profile $\delta F(\vec{\gamma}_i, t+\tau_i)$, $k$ is the emitter index ($k$ goes from $k$=0 to $k$=$N$ where $N$ is the total number of emitters in the sample), $\epsilon_k$ is the 'on'-state brightness of the $k^{\text{th}}$ emitter, $\delta b_k(t+\tau_i)$ is the blinking profile fluctuation of the $k^{\text{th}}$ emitter with time shift $\tau_i$, and $U(\vec{r})$ is the PSF of the optical system. It is approximated with a Gaussian function (equation (2.11), main text). The expression $G(I_p)$ with $I_p = \{\delta F(\vec{\gamma}_i, t+\tau_o) \,|\, i \in [1,m]\}$ is given by:

$$G(I_p) = \left\langle \prod_{i=1}^{m} \delta F(\vec{r}_i, t+\tau_i) \right\rangle_t \qquad \text{(SI-1.3)}$$

By substituting the expression for $\delta F(\vec{r}_i, t)$ shown in equation (SI-1.2) into equation (SI-1.3), we get:

$$\begin{aligned} G(I_p) &= \left\langle \prod_{i=1}^{m} \left( \sum_{k=1}^{N} \epsilon_k \cdot \delta b_k(t+\tau_i)\cdot U(\vec{\gamma}_i - \vec{r}_k) \right) \right\rangle_t \\ &= \sum_{k_1=1}^{N}\sum_{k_2=1}^{N}\cdots\sum_{k_m=1}^{N} \left( \prod_{i=1}^{m} \epsilon_{k_i} \right) \cdot \left\langle \delta b_{k_1}(t+\tau_1)\cdots \delta b_{k_m}(t+\tau_m) \right\rangle_t \cdot \left( \prod_{i=1}^{m} U(\vec{\gamma}_i - \vec{r}_{k_i}) \right) \end{aligned} \quad , \qquad \text{(SI-1.4)}$$

where $m$ is the total number of elements in $I_p$.

When we express a certain partition of set $I$ as $\{I_1, I_2, ..., I_q\}$, where $I_1 \cup I_2 \cup \cdots \cup I_q = I$, the products of all the cross-correlation functions of the subset need to be analyzed:

$$G(I_1)\cdot G(I_2)\cdots G(I_q) \qquad \text{(SI-1.5)}$$

Note that $q$ is the total number of parts in this partition. The number of terms in the multiplication of all cross-correlation functions of all the different parts ( total number of $q$ subsets) will always equal to $n$, (being equal to the size of $I$ ). This means that when substituting equation (SI-1.4) into (SI-1.5), the resulted expression will always yield a $n$-fold summation series. We can use a set of $n$ indexes $\{k_1,k_2,\ldots,k_n\}$ to denote the emitter indexes for the $n$-fold summation series. Note that in the expanded expression after substituting equation (SI-1.4) into (SI-1.5) with different partitions of $I$, the differences would result only in the factor that involves averages over time: $<\delta b_{k1}(t+t_1)\cdot \delta b_{k2}(t+t_2)\cdot\ldots\cdot \delta b_{km}(t+t_m)>_t$, because the set of indexes $\{k_1,k_2,\ldots,k_m\}$ in (SI-1.4) changes with the change of partitions. This is because the partitions of $\{k_1,k_2,\ldots,k_n\}$ and $\{\delta b_{k1}(t+\tau_1), \delta b_{k2}(t+\tau_2),\ldots,\delta b_{kn}(t+\tau_n)\}$ are both defined by the same partitions of the set $I$ , because both are partitioned by the same set of indexes.

Let's denote $B$ as this set (i.e. B={ $\delta b_{k1}(t+\tau_1), \delta b_{k2}(t+\tau_2),\ldots,\delta b_{kn}(t+\tau_n)\}$), with a certain partition of $B$ expressed as: $\{B_1, B_2,\ldots, B_q\}$ such that $B_1 \cup B_2 \cdots \cup B_q = B$ . This partition is also associated with a partition of set $I$ expressed as $\{I_1, I_2,\ldots, I_q\}$ such that $I_1 \cup I_2 \cup \cdots \cup I_q = I$ . The cross-cumulant of set $B_p$, denoted by $G(B_p)$, can then be written:

$$\begin{aligned} &G(B_p) = \left\langle \delta b_{k_1}(t+\tau_1)\cdot \delta b_{k_2}(t+\tau_2)\cdots \delta b_{k_m}(t+\tau_m)\right\rangle_t , \\ &\text{with} \quad B_p = \{\delta b_{k_1}(t+\tau_1), \delta b_{k2}(t+\tau_2),..., \delta b_{k_m}(t+\tau_m)\} \end{aligned} \qquad \text{(SI-1.6)}$$

Combining (SI-1.4) and (SI-1.6) into (SI-1.1) , we get:

$$C_n(I) = \sum_{\substack{\text{all partitions}\\ \text{of } I}} \left\{ \left[(-1)^{q-1}(q-1)!\right] \cdot \left[ \sum_{k_1=1}^{N} \cdots \sum_{k_n=1}^{N} \left(\prod_{i=1}^{n} \epsilon_{k_i}\right) \cdot \left(\prod_{p=1}^{q} G(B_p)\right) \left(\prod_{i=1}^{n} U(\vec{\gamma}_i - \vec{r}_{k_n})\right) \right] \right\} \qquad \text{(SI-1.7)}$$

Note here that $\{B_1, B_2,\ldots, B_q\}$ is the partition of $B$, that corresponds to $\{I_1,I_2,\ldots, I_q\}$ as partitions of $I$ that is partitioned by the same group of indexes. Because only the factor in the inner most summation series that is partition-dependent is the serial product of $G(B_p)$ corresponding to different partitions, we can therefore change the order of the $n$-fold summation series to the outer layer of the summation series, together with the other two factors other than the factor involves $G(B_p)$. We therefore get:

$$C_n(I) = \left[ \sum_{k_1=1}^{N} \cdots \sum_{k_n=1}^{N} \left(\prod_{i=1}^{n} \epsilon_{k_i}\right) \cdot \left\{ \sum_{\substack{\text{all partitions}\\ \text{of } B=\{\delta b_1,\ldots,\delta b_n\}}}^{B_1\cup\cdots\cup B_q = B} \left[(-1)^{q-1}(q-1)!\right] \cdot \prod_{p=1}^{q} G(B_p) \right\} \left(\prod_{i=1}^{n} U(\vec{\gamma}_i - \vec{r}_{k_n})\right) \right] \qquad \text{(SI-1.8)}$$

Notice the similarity between the form in the curly parentheses and equation (SI-1.1). This form is in fact the cross-cumulants of set ***B***:

$$C_n(B) = \sum_{\substack{\text{all partitions}\\ \text{of } B=\{\delta b_1,\ldots,\delta b_n\}}}^{B_1\cup\cdots\cup B_q = B} \left[(-1)^{q-1}(q-1)!\right] \cdot \prod_{p=1}^{q} G(B_p) \qquad \text{(SI-1.9)}$$

Since cross-cumulants vanish to zero when a subset is independent of the remainder[2, 3], the cross-cumulant would be zero for the set $B = \{\delta b_{k_1}(t+\tau_1), \delta b_{k2}(t+\tau_2),..., \delta b_{k_n}(t+\tau_n)\}$ where the indexes $\{k_1,k_2,\ldots,k_n\}$ are all identical because we assume the emitters blinking independently. We can therefore rephrase this statement:

$$C_n(B) = \begin{cases} \text{non-zero;} & \text{when } k_1 = k_2 = \cdots = k_n \\ 0; & \text{otherwise} \end{cases} \qquad \text{(SI-1.10)}$$

and drop the zero terms from equation (SI-1.8) to yield a simplified form (equation SI-1.11):

$$C_n(I) = \sum_{k=1}^{N} \epsilon_k^n \cdot C_n(\delta b_{k_1}(t+\tau_1), \delta b_{k2}(t+\tau_2),..., \delta b_{k_n}(t+\tau_n)) \left(\prod_{i=1}^{n} U(\vec{\gamma}_i - \vec{r}_k)\right) \qquad \text{(SI-1.11)}$$

Note that the non-zero terms have $k_1=k_2=\cdots=k$. We also simplified $\epsilon_{k1}\cdot\epsilon_{k2}\cdots\epsilon_{k3}$ into $\epsilon_k^n$ in equation (SI-1.11) above.

Next, we use the relation (SI-1.12) (a proof is given at the bottom of this note):

$$\begin{aligned} &\prod_{i=1}^{n} U(\vec{\gamma}_i - \vec{r}_k) = W_n(\vec{\gamma}_1,..,\vec{\gamma}_n) U^n(\vec{\gamma}_{gc} - \vec{r}_k) \\ &\text{with} \quad \vec{\gamma}_{gc} = \frac{1}{n}\sum_{k=1}^{n} \vec{\gamma}_k \ ; \quad W_n(\vec{\gamma}_1,...,\vec{\gamma}_n) = \prod_{\substack{i\in[1,n]\\ j\in[1,n]}}^{i<j} U\left(\frac{\vec{\gamma}_i - \vec{\gamma}_j}{\sqrt{n}}\right) ; \end{aligned} \quad , \qquad \text{(SI-1.12)}$$

and Gaussian approximation of $U(\vec{r})$: $\quad U(\vec{r}) = \exp\left(-\frac{r_x^2 + r_y^2}{2\omega_{xy}^2} - \frac{r_z^2}{2\omega_z^2}\right)$

where $\vec{\gamma}_i$ represents the location vector of pixel *i* (to contrast with location vector of emitter k, $\vec{r}_k$) and n is the total number of elements in set I (also the cumulant order). Substituting (SI-1.12) into (SI-1.11) yields:

$$C_n(I) = \sum_{k=1}^{N} \epsilon_k^n C_n\left(\delta b_k(t+\tau_1), \delta b_k(t+\tau_2), ..., \delta b_k(t+\tau_n)\right) \cdot W_n(\vec{r}_1, \vec{r}_2, ..., \vec{r}_n) \cdot U^n(\vec{r}_{gc} - \vec{r}_k) \quad . \qquad \text{(SI-1.13)}$$

Since $W_n(\vec{r}_1, \vec{r}_2, ..., \vec{r}_n)$ is independent of emitter index $k$, it can be taken out of the summation series:

$$\begin{aligned} & C_n(I) \\ & = W_n(\vec{r}_1, \vec{r}_2, ..., \vec{r}_n) \cdot \sum_{k=1}^{N} \epsilon_k^n C_n\left(\delta b_k(t+\tau_1), \delta b_k(t+\tau_2), ..., \delta b_k(t+\tau_n)\right) \cdot U^n(\vec{r}_{gc} - \vec{r}_k) \\ & \quad \text{with:} \\ & \quad W(\vec{r}_1, \vec{r}_2, ..., \vec{r}_n) = \prod_{\substack{i \in [1,n] \\ j \in [1,n]}}^{i<j} U\left(\frac{\vec{r}_i - \vec{r}_j}{\sqrt{n}}\right) \end{aligned} \qquad \text{(SI-1.14)}$$

We hence derived equation 2.6 of the main text.

**Proof of equation (SI-1.12):**

$$\begin{aligned} & \textit{to prove}: \\ & \prod_{i=1}^{n} U(\vec{\gamma}_i - \vec{r}_k) = W_n(\vec{\gamma}_1, .., \vec{\gamma}_n) U^n(\vec{\gamma}_{gc} - \vec{r}_k) \\ & \text{with} \quad \vec{\gamma}_{gc} = \frac{1}{n}\sum_{k=1}^{n} \vec{\gamma}_k \quad ; \quad W_n(\vec{\gamma}_1, ..., \vec{\gamma}_n) = \prod_{\substack{i \in [1,n] \\ j \in [1,n]}}^{i<j} U\left(\frac{\vec{\gamma}_i - \vec{\gamma}_j}{\sqrt{n}}\right); \\ & \text{and Gaussian approximation of } U(\vec{r}): \quad U(\vec{r}) = \exp\left(-\frac{r_x^2 + r_y^2}{2\omega_{xy}^2} - \frac{r_z^2}{2\omega_z^2}\right) \end{aligned}$$

In order to prove (SI-1.12), we divide both sides of the equation by $U^n(\vec{\gamma}_{gc} - \vec{r}_k)$ and substitute in the (Gaussian) analytical expression for $W_n(\vec{\gamma}_1, ..., \vec{\gamma}_n)$ :

$$\frac{\prod_{i=1}^{n} U(\vec{\gamma}_i - \vec{r}_k)}{U^n(\vec{\gamma}_{gc} - \vec{r}_k)} = \prod_{\substack{i \in [1,n] \\ j \in [1,n]}}^{i<j} U\left(\frac{\vec{\gamma}_i - \vec{\gamma}_j}{\sqrt{n}}\right) \qquad \text{(SI-1.15)}$$

the left-hand side of equation (SI-1.15) can therefore be written as:

$$\begin{aligned} & \frac{\prod_{i=1}^{n} U(\vec{\gamma}_i - \vec{r}_k)}{U^n(\vec{\gamma}_{gc} - \vec{r}_k)} \\ & = \exp\left[-\sum_{i=1}^{n}\left(\frac{(\gamma_{i,x} - r_{k,x})^2 + (\gamma_{i,y} - r_{k,y})^2}{2\omega_{xy}^2} + \frac{(\gamma_{i,z} - r_{k,z})^2}{2\omega_z^2}\right)\right] \\ & \cdot \exp\left[-n \cdot \left(\frac{(\gamma_{gc,x} - r_{k,x})^2 + (\gamma_{gc,y} - r_{k,y})^2}{2\omega_{xy}^2} + \frac{(\gamma_{gc,z} - r_{k,z})^2}{2\omega_z^2}\right)\right] \\ & = \prod_{d=\{x,y,z\}} \left\{ \exp\left[-\frac{1}{2\omega_d^2}\left(\sum_{i=1}^{n}(\gamma_{i,d} - r_{k,d})^2 - n(\gamma_{gc,d} - r_{k,d})^2\right)\right]\right\} \\ & \text{with } \omega_x = \omega_y = \omega_{xy}; \quad \textit{and} \ \gamma_{gc,d} = \frac{1}{n}\sum_{i=1}^{n} \gamma_{i,d} \end{aligned} \qquad \text{(SI-1.16)}$$

Next, we substitute the Gaussian approximation into the right-hand side of (SI-1.15) to get:

$$\prod_{\substack{i\in[1,n]\\ j\in[1,n]}}^{i<j} U\left(\frac{\vec{\gamma}_i-\vec{\gamma}_j}{\sqrt{n}}\right)$$

$$=\exp\left[-\sum_{\substack{i\in[1,n]\\ j\in[1,n]}}^{i<j}\left(\frac{(\gamma_{i,x}-\gamma_{j,x})^2+(\gamma_{i,y}-\gamma_{j,y})^2}{2\omega_{xy}^2\cdot n}+\frac{(\gamma_{i,z}-\gamma_{j,z})^2}{2\omega_z^2\cdot n}\right)\right] \qquad \text{(SI-1.17)}$$

$$==\prod_{d=\{x,y,z\}}\left\{\exp\left[-\frac{1}{2\omega_d^2}\left(\frac{1}{n}\cdot\sum_{i=1}^{n}(\gamma_{i,d}-\gamma_{j,d})^2\right)\right]\right\}$$

$$\text{with } \omega_x=\omega_y=\omega_{xy}\,; \quad \text{and } \gamma_{gc,d}=\frac{1}{n}\sum_{i=1}^{n}\gamma_{i,d}$$

Proving (SI-1.15) is equivalent to proving the equality between (SI-1.16) and (SI-1.17), which is equivalent to prove the following relation:

$$\sum_{i=1}^{n}(\gamma_{i,d}-r_{k,d})^2-n(\gamma_{gc,d}-r_{k,d})^2=\frac{1}{n}\cdot\sum_{i,j\in[1,n]}^{i<j}(\gamma_{i,d}-\gamma_{j,d})^2 \qquad \text{(SI-1.18)}$$

By substituting $\gamma_{gc,d}=\frac{1}{n}\sum_{i=1}^{n}\gamma_{i,d}$ (equation SI-1.16) into (SI-1.18), and dropping the indices $d$ and $k$ (to simplify the notation), we get:

$$\sum_{i=1}^{n}(\gamma_i-r)^2-n\cdot\left(\left(\frac{1}{n}\sum_{i=1}^{n}\gamma_i\right)-r\right)^2=\frac{1}{n}\cdot\sum_{i,j\in[1,n]}^{i<j}(\gamma_i-\gamma_j)^2 \qquad \text{(SI-1.19)}$$

We next expand and simplify the two terms on the left side of (SI-1.19) to yield:

$$\sum_{i=1}^{n}(\gamma_i-r)^2-n\cdot\left(\left(\frac{1}{n}\sum_{i=1}^{n}\gamma_i\right)-r\right)^2$$

$$=\sum_{i=1}^{n}(\gamma_i^2-2\cdot r\cdot\gamma_i+r^2)-n\cdot\left(\left(\frac{1}{n}\sum_{i=1}^{n}\gamma_i\right)^2-2r\cdot\left(\frac{1}{n}\sum_{i=1}^{n}\gamma_i\right)+r^2\right)$$

$$=\sum_{i=1}^{n}(\gamma_i^2-2\cdot r\cdot\gamma_i+r^2)-\left(\frac{1}{n}\left(\sum_{i=1}^{n}\gamma_i\right)^2-2r\cdot\sum_{i=1}^{n}\gamma_i+n\cdot r^2\right) \qquad \text{(SI-1.20)}$$

$$=\sum_{i=1}^{n}\gamma_i^2-2\cdot r\sum_{i=1}^{n}\gamma_i+n\cdot r^2-\frac{1}{n}\left(\sum_{i=1}^{n}\gamma_i\right)^2+2r\cdot\sum_{i=1}^{n}\gamma_i-n\cdot r^2$$

$$=\sum_{i=1}^{n}\gamma_i^2-\frac{1}{n}\left(\sum_{i=1}^{n}\gamma_i\right)^2$$

Now we only need to prove that this simplified expression equals to the right side of (SI-1.19):

$$\sum_{i=1}^{n}\gamma_i^2-\frac{1}{n}\left(\sum_{i=1}^{n}\gamma_i\right)^2=\frac{1}{n}\cdot\sum_{i,j\in[1,n]}^{i<j}(\gamma_i-\gamma_j)^2\ . \qquad \text{(SI-1.21)}$$

Let's multiply both sides of equation (SI-1.21) by $2\cdot n$ :

$$2 \cdot n \cdot \sum_{i=1}^{n} \gamma_i^2 - 2\,(\sum_{i=1}^{n} \gamma_i)^2 = 2 \sum_{i,j \in [1,n]}^{i<j} (\gamma_i - \gamma_j)^2 \qquad \text{(SI-1.22)}$$

but since:

$$2 \cdot n \cdot \sum_{i=1}^{n} \gamma_i^2 = 2 \cdot \sum_{i=1}^{n}\sum_{j=1}^{n} \gamma_i^2 = \sum_{i=1}^{n}\sum_{j=1}^{n} \left(\gamma_i^2 + \gamma_j^2\right) \qquad \text{(SI-1.23)}$$

we can substitute (SI-1.23) into the left hand side of (SI-1.22) to get:

$$\begin{aligned}
&2 \cdot n \cdot \sum_{i=1}^{n} \gamma_i^2 - 2\,(\sum_{i=1}^{n} \gamma_i)^2 \\
&= \sum_{i=1}^{n}\sum_{j=1}^{n} \left(\gamma_i^2 + \gamma_j^2\right) - 2\,(\sum_{i=1}^{n} \gamma_i)^2 \\
&= \sum_{i=1}^{n}\sum_{j=1}^{n} \left(\gamma_i^2 + \gamma_j^2\right) - \sum_{i=1}^{n}\sum_{j=1}^{n} 2\gamma_i\gamma_j = \sum_{i=1}^{n}\sum_{j=1}^{n} \left(\gamma_i^2 + \gamma_j^2 - 2\gamma_i\gamma_j\right) \\
&= \sum_{i=1}^{n}\sum_{j=1}^{n} (\gamma_i - \gamma_j)^2
\end{aligned} \qquad \text{(SI-1.24)}$$

We then substitute (SI-1.24) into (SI-1.22):

$$\sum_{i=1}^{n}\sum_{j=1}^{n} (\gamma_i - \gamma_j)^2 = 2 \sum_{i,j \in [1,n]}^{i<j} (\gamma_i - \gamma_j)^2 \qquad \text{(SI-1.25)}$$

Because of index symmetry,

$$\sum_{i,j \in [1,n]}^{i<j} (\gamma_i - \gamma_j)^2 = \sum_{i,j \in [1,n]}^{i>j} (\gamma_i - \gamma_j)^2 \qquad \text{(SI-1.26)}$$

As a result, the right hand side of (SI-1.25) becomes:

$$2 \sum_{i,j \in [1,n]}^{i<j} (\gamma_i - \gamma_j)^2 = \sum_{i,j \in [1,n]}^{i<j} (\gamma_i - \gamma_j)^2 + \sum_{i,j \in [1,n]}^{i>j} (\gamma_i - \gamma_j)^2 \qquad \text{(SI-1.27)}$$

Additionally, since:

$$\sum_{i,j \in [1,n]}^{i=j} (\gamma_i - \gamma_j)^2 = 0 \ , \qquad \text{(SI-1.28)}$$

we can further simplify the right hand side of equation (SI-1.25) :

$$\begin{aligned}
&2 \sum_{i,j \in [1,n]}^{i<j} (\gamma_i - \gamma_j)^2 \\
&= \sum_{i,j \in [1,n]}^{i<j} (\gamma_i - \gamma_j)^2 + \sum_{i,j \in [1,n]}^{i>j} (\gamma_i - \gamma_j)^2 + \sum_{i,j \in [1,n]}^{i=j} (\gamma_i - \gamma_j)^2 \\
&= \sum_{i,j \in [1,n]}^{i<j,\, i=j,\, i>j} (\gamma_i - \gamma_j)^2 \\
&= \sum_{i=1}^{n}\sum_{j=1}^{n} (\gamma_i - \gamma_j)^2
\end{aligned} \qquad \text{(SI-1.29)}$$

Which is identical to the left hand side of equation (SI-1.25).

We have thus proved the identity (SI-1.25) and hence the identity(SI-1.12).

### Appendix 2

This note discusses a strategy for the selection of pixel combination for high order cumulants to yield a virtual pixel located at a specific sub grid as compared to the grids of the original pixel size.

Every SOFI pixel in a $n^{th}$ order SOFI image is calculated as the joint cumulant of a set of n elements (in the form of pixels with or without duplicated pixels) selected from a supply of real detector pixels. In Fig. S2, we demonstrated the case of 5th order SOFI to explain how to choose the pixel combinations. We demonstrated a pixel array with four real detector pixels (P1, P2, P3, P4) as the supply (Fig. S2(a)), and all possible virtual pixel locations corresponding to 5th order. Each virtual pixel [4] requires the 5-element set (A, B, C, D, E) constructed with elements from the real pixel supply (P1, P2, P3, P4), with the allowance of duplicated pixels. For a given SOFI pixel location, the choice of the five elements from the real pixel supply (P1, P2, P3, P4) determines the geometrical center of the finer-grid SOFI pixel. The total number of possible pixel combinations is astronomical. In the example of SI Fig. S2(a), the number of possible pixel combinations is 45=1024. Obviously, if the supply of real pixels is increased, the possible pixel combinations is increased even further.

An efficient way to search for such pixel combinations can be guided by a geometric interpretation of the geometry center of vectors (Fig. S2(b) and S2(c)). We can define location vectors of each pixel in the set (A, B, C, D, E) as $\{\vec{r}_A, \vec{r}_B, \vec{r}_C, \vec{r}_D, \vec{r}_E\}$ respectively. Then the geometry center of these five real pixel locations would be $\vec{r}_{gc} = (\vec{r}_A + \vec{r}_B + \vec{r}_C + \vec{r}_D + \vec{r}_E)/5$. If we denote $\vec{r}_x' = \vec{r}_x / 5$ where $x$ represents A, B, C, D, and E, the calculation of the geometry center $\vec{r}_{gc}$ is equivalent to aligning all the vectors of set $\{\vec{r}_x'\}$ one after another (with any order) from any pre-defined origin point (Fig. S2 (b)). As a result, the ending point of the last vector would be the location pointed to by vector $\vec{r}_{gc}$.

In our example (Fig. S2), each vector $\vec{r}_x$ and $\vec{r}_x'$ can have 4 different possibilities based on the choices of real pixels. Also, the vectors are denoted as $\vec{r}_i$ and $\vec{r}_i'$ where $i$ = 1, 2, 3, 4. We define the origin point at the location of P3. The direction vectors of real pixels as shown at the bottom of Fig. S2(a). Note here that based on the definition of origin (at the location of P3), $\vec{r}_3 = \vec{0}$ (so both $\vec{r}_3$ and $\vec{r}_3'$ do not have directions). The length of one location vector (with $\vec{r}_4$ shown as an example in Fig. S2(a)) is the distance from the origin point (P3 in this example) to the pixel. The length of $\vec{r}_i'$ is one fifth of the original location vector (see $\vec{r}_4'$ as an example in Fig. S2(a)). The geometric summations of the vectors $\{\vec{r}_i'\}$ are shown in Fig. S2(b) and (c), where two possible choices of pixel combinations are demonstrated to show that both yield virtual pixel V1 as labeled in Fig. S2(a), (b), (c). Based on the vectors used in the alignment, we can get the corresponding pixel combinations as shown in Fig. S2(d) and (e) respectively; in Fig. S2(d) the set of five pixels is (P2, P3, P3, P4, P4), and in Fig. S2(e) the set is (P1, P3, P3, P3, P4). Note that the repeating pixel P3 adds a vector of length=0, so the repeating $\vec{r}_3'$ vectors are overlapping (2 or 3 filled blue circles in Fig. S2 (b) and S2(c) respectively).

### Appendix 3

The derivation of the analytical form for the reconstructed moments $M_n(\vec{r})$ (equation (2.14) of main text) is given below. This is composed of terms containing information on the PSF of the optical system, and the locations, brightness and blinking profile of all the emitters in the field-of-view.

We start with the expression for the fluorescence fluctuation (equation (2.5) in the main text). With the extension of finer-grid and the re-mapping of the cumulant image, the accessible locations of cumulants are expanded from the limited locations of SOFI pixels to the finer-grid pixels. The SOFI cumulants of these fluctuations can be written as:

$$C_n(\vec{r}) = C_n(\delta F(\vec{r},t)) = C_n\left(\sum_{k=1}^{N}\epsilon_k \cdot \delta b_k(t) \cdot U(\vec{r}-\vec{r}_k)\right) \quad . \qquad \text{(SI-3.1)}$$

Because cumulants are additive, and only $\delta b_k(t)$ is time-dependent, we can change the order in the summation:

$$C_n(\vec{r}) = \sum_{k=1}^{N}\epsilon_k^n C_n(\delta b_k(t)) U^n(\vec{r}-\vec{r}_k) \quad . \qquad \text{(SI-3.2)}$$

A single term in the summation series can be expressed as:

$$\begin{aligned} C_{s_1}(\vec{r})\cdots C_{s_\sigma}(\vec{r}) &= \left(\sum_{k=1}^{N}\epsilon_k^{s_1} C_{s_1}(\delta b_k(t)) U^{s_1}(\vec{r}-\vec{r}_k)\right)\cdots\left(\sum_{k=1}^{N}\epsilon_k^{s_\sigma} C_{s_\sigma}(\delta b_k(t)) U^{s_\sigma}(\vec{r}-\vec{r}_k)\right) \\ &= \sum_{k_1=1}^{N}\cdots\sum_{k_\sigma=1}^{N}\left(\prod_{p=1}^{\sigma}\epsilon_{k_p}^{s_p}\right)\left(\prod_{p=1}^{\sigma}C_{s_p}(\delta b_{k_p}(t))\right)\left(\prod_{p=1}^{\sigma}U^{s_p}(\vec{r}-\vec{r}_{k_p})\right) \end{aligned} \qquad \text{(SI-3.3)}$$

Where $\{I_p \mid p = 1,2,...,\sigma\}$ denotes the term contributed by a specific partition of set $\{1,2,...,n\}$, and $s_p$ denotes the size of part $I_p$. By re-arranging the third factor in the summation series from equation SI-3.3 in a similar way to the derivation of equation (2.6) from the main text and expanded in Appendix 1) we get:

$$\prod_{p=1}^{\sigma}U^{s_p}(\vec{r}-\vec{r}_{k_p}) = U^n(\vec{r}-\frac{1}{n}\sum_{p=1}^{\sigma}s_p\cdot\vec{r}_{k_p})W(\{s_p,\vec{r}_{k_p}\}) \quad \text{with } \mathrm{p}\in[1,\sigma] \ , \qquad \text{(SI-3.4)}$$

where we dub $W$ as the “emitter distance factor”, whose analytical form is identical to that of the distance factor introduced in previous work [5] (see equation (SI-1.12)). The emitter distance factor is determined by the set of emitter location vectors defined by $\{(s_p,\vec{r}_{k_p}) \mid p\in[1,\sigma]\}$ such that each $\vec{r}_{k_p}$ is repeated $s_p$ times to form a set with a total of $\sum_{p=1}^{\sigma}s_\sigma$ elements. Substituting equation (SI-3.4) into equation (SI-3.3) yields:

$$C_{s_1}(\vec{r})\cdots C_{s_\sigma}(\vec{r}) = \sum_{k_1=1}^{N}\cdots\sum_{k_\sigma=1}^{N}\left(\prod_{p=1}^{\sigma}\epsilon_{k_p}^{s_p}\right)\left(\prod_{p=1}^{\sigma}C_{s_p}(\delta b_{k_p}(t))\right)U^n(\vec{r}-\frac{1}{n}\sum_{p=1}^{\sigma}s_p\cdot\vec{r}_{k_p})W(\{s_p,\vec{r}_{k_p}\}) \qquad \text{(SI-3.5)}$$

Lastly, by substituting equation (SI-3.5) into equation (3.3) from the main text, we derive the analytical form for moments reconstructed moments:

$$\begin{aligned} M_n(\vec{r};\tau=0) &= \sum_{\substack{I_1\cup I_2\cup\cdots\cup I_\sigma=\{1,2,...,n\} \\ \text{all partitions of set }\{1,2,...,n\}}} C_{s_1}(\vec{r},\tau=0)\cdots C_{s_\sigma}(\vec{r},\tau=0) \\ &= \sum_{\substack{I_1\cup I_2\cup\cdots\cup I_\sigma=\{1,2,...,n\} \\ \text{all partitions of set }\{1,2,...,n\}}}\left[\sum_{k_1=1}^{N}\cdots\sum_{k_\sigma=1}^{N}\left(\prod_{p=1}^{\sigma}\epsilon_{k_p}^{s_p}\right)\left(\prod_{p=1}^{\sigma}C_{s_p}(\delta b_{k_p}(t))\right)U^n(\vec{r}-\vec{r}_m)W(\{s_p,\vec{r}_{k_p}\})\right] \end{aligned} \qquad \text{(SI-3.6)}$$

where $\vec{r}_m$ is defined as

$$\vec{r}_m = \frac{1}{n}\sum_{p=1}^{\sigma}s_p\cdot\vec{r}_{k_p} \qquad \text{(SI-3.7)}$$

### Appendix 4

We discuss here the local dynamic range compression (*ldrc*) algorithm.

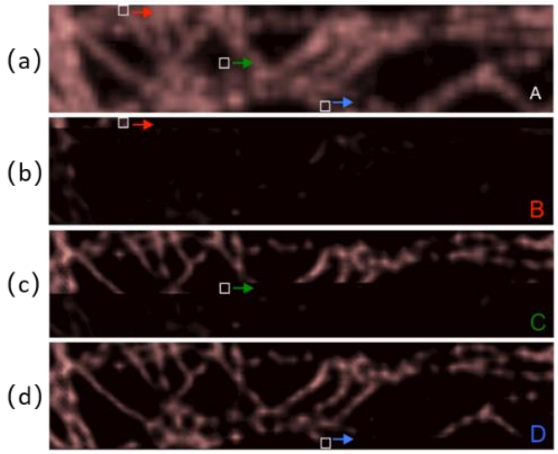

Fig. S16. Local dynamic range compression. Panel (a) is the reference image, where a small window is defined in the image as shown in the figure above. This window slides pixel-wise across the field-of-view as shown in (b), (c) and (d) (color coded to show different window locations shown in A). The computation can be implemented in a parallel manner.

High order cumulants or moments reconstructions result-in images with a large dynamic range of pixel intensities. The ldrc algorithm compresses the dynamic range of these reconstructions with respect to a reference image (second order SOFI image or the average/sum image) while retaining resolution enhancement. By way of example, Fig. S16 demonstrates the algorithm. Fig. S16(a) shows the reference (in this case the average image), while Figs. S16(b) – (d) show the progression of algorithm across the image. Dynamic range compression is performed locally in a small window that is scanned across the image. It can be performed either by faster-scanning a single window across the whole image or by scanning several windows in parallel. Fig. S16 demonstrates the implementation of ldrc with a single sliding window. The dynamic range of pixel intensities within the windowed area is linearly rescaled with respect to the dynamic range of the pixels of the same window in the reference image. As explained below:

$$I_i^* = \frac{I_i - I_l}{I_u - I_l} \cdot R_u \qquad \text{(SI-4.1)}$$

Where $I_i$ is the $i^{th}$ pixel in the window from the high dynamic range input image. $I_l$ and $I_u$ are the lower and upper bound of the pixel intensities in the window from the high dynamic range input image. And $R_u$ is the upper bound (maximum value) of the pixel intensities in the window of the reference image, $I_i^*$ is the pixel value after this linear rescale of the specific window.

Fig. S16(b) – (d) show intermediate results during the progression of the algorithm, sampled at window positions progressively from left to right, from top to bottom up to the position marked on the corresponding panels. Only one window is used to simplify the illustration. Note that because the window contains multiple pixels and slides pixel-wise, therefore each individual pixel will appear in multiple windows, yielding multiple rescaled pixel intensities $I_i^*$ that correspond to multiple windows. The final output pixel intensity is averaged across all the yielded $I_i^*$ from multiple windows. This averaging, however, is somewhat reduced near the image edges (as the pixels are covered by less number of windows). The ldrc code (xy_QuickLDRC.m) is posted on Github as part of the SOFI2.0 pacakge [6].

### Appendix 5

This note details the simulations used for this work. We start in **5.1** with a general description of the simulations developed with a modular design. Specifics of each module used in this study are discussed in **5.2**. Details of each specific simulation dataset are discussed in **5.3**.

The simulation package was developed in MATLAB and is an advanced version of the simulation developed for our previous work [7]. The modularized datasets were stored in the form of *.mat files. The source code package is posted on Github as ***SR_Simu3D***[8].

#### *5.1. General description of the simulator*

Here we present the general description of the algorithm used in the simulator, and is explained in Chart 1 (see below). The simulator consists of 6 major steps, where the functionality of each step is designed in the form of a module that are independent and replaceable. The sequence of the modules resembles the natural process of first preparing the sample, then forming the image based on the optical system used, and finally acquiring the data. The input of each module could either be the output from the upper-stream module, or loaded from pre-computed results from the upper-stream module. The output of each module could either be pre-computed or computed on demand.

#### Chart 1. Architecture of the simulator

**1.** Get a labeled sample in 3D.
- **1.1.** Get feature of interest with close-packed available labeling sites.
- **1.2.** Get the selected labeling site
- **1.3.** Get the emitter locations considering labeling uncertainty.

**2.** Get time-intensity trajectories for each individual emitter.
- **2.1** Get the blinking sequence in terms of on/off time durations.
- **2.2** Get the blinking sequence in terms of photon numbers per frame.

**3.** Get time-location trajectories for each individual emitter. (don’t use if feature is static)

**4.** Get PSF model (in the form of a 3D matrix). (independent from step 1 to 3)

<u>*Loop 1. For all frames*</u> (or compute in parallel)

**5.** Generate images formed at the detection plane of the simulated sample.

<u>*Loop 2. For all emitters*</u> (or compute in parallel)
- **5.1.** Determine the PSF of this emitter in detection plane using the emitter location and the precomputed 3D PSF matrix from step 4.
- **5.2.** Calculate the arrival locations of the photons emitted by this emitter in the detection plane, where the probability distribution function of the arrival locations is described by the image determined in step 5.1.

<u>*End of loop 2.*</u>

**6.** Generate images captured by the detector array (each detector is a pixel).
- **6.1.** Calculate the number of photons arrived at each pixel.
- **6.2.** Calculate the detected number of photons at each pixel.
  - **6.3.** Add the background signal/noise to each pixel.
  - **6.4.** Export the calculated frame. (.tif format)

<u>*End of Loop 1*</u>

Below we describe the overall modular architecture of the simulator, the interfaces for each module, and the elements required (or provided) for each module. The first step constitutes input parameters that specify the simulated sample (such as sample volume, labeling density labeling uncertainty, grid size, etc.) to create a labeled sample in 3D volume. Specifically, in step 1.1 we generate features of interest (sample morphology such as filaments, vesicles, aggregates, etc.), where an array of x-y-z coordinates characterizes the available labeling sites. In step 1.2, a second array of x-y-z coordinates that were randomly selected from the first array is generated (representing a sub-set from all available sites as labeled sites). Labeling uncertainties are simulated subsequently in step 1.3 by updating the x-y-z coordinates of the labeled sites with random errors based on a Gaussian distribution with a user-defined width, resulting in a more-realistic third array of emitter locations (i.e. x-y-z coordinates).

In the second step we calculate the blinking sequences of each emitter. In step 2.1, on/off duration times are calculated for each emitter (with a double float precision, non-digitalized). Subsequently, a blinking trajectory for each emitter (as a 1D array) is generated (based on a user-defined integration time per frame). The $i^{th}$ element in this array represents the percentage of time the emitter is in the 'on' state during the $i^{th}$ frame. In step 2.3 the time trajectory 1D array is multiplied by a scalar representing a predefined photon budget. The resulted 1D array represents the total number of photons per time-bin for a given emitter (the $i^{th}$ element contains the number of photons detected during the $i^{th}$ frame or $i^{th}$ time-bin).

To account for feature dynamics (representing dynamic morphological changes in live cells) the third step calculates the time-location trajectory for each emitter (the location is constant for static features). Time-varying emitter location coordinates can simulate instrument drift, diffusion, or directed motion (as in the case of cell motility) or their combination, with the possibility for further extensions.

The fourth step is independent from the first 3 steps and it accounts for the 3D point-spread-function (PSF) of the optical system represented as a 'PSF matrix'. The center of the PSF is located at the center of the 3D matrix. This step can be performed independently and in parallel. Alternatively, a precomputed PSF matrix library could be called by the algorithm.

The following steps 5 and 6 operate on individual frames of the simulated movie. In the fifth step, a 2D matrix representation of the sample's image at the detection plane is calculated. This image is the super-position of images formed by all the emitters in the simulated sample, and the image of each individual emitter is determined by four factors: The 3D PSF matrix, the total number of photons emitted by each emitter, the x-y-z coordinates of the emitter, and the defined z coordinate of the focal plane. The point spread function of each emitter is determined in step 5.1, and the actual arrival locations of the photons emitted by this emitter within the given frame is determined in step 5.2. In this step, we generate arrival locations of photons emitted by individual emitters. First the PSF image of this emitter is determined that will describe the 2D probability distribution of the arrival locations of the photons emitted by this emitter within this time-bin, and the total number of photon emitted by this emitter is pre-calculated with the time-intensity trajectories. Once the photon arrival locations for all the emitters are determined, they will constitute another 2D matrix that represent the full field of view of the simulated imaging system with all the arrived photons positioned in the matrix. Each element of this matrix represents the given location in the field of view, and the value of this element represents the number of photons that have arrived in the corresponding area.

In the sixth step, we calculate the image captured by the detector array of the camera. This step operates on the movie frame calculated in step 5 to bin the data onto the predefined pixel grid of the detector (according to the pre-defined / user selected pixel size). Step 6.1 simulates the binning effect per detector area. The total number of photons per emitter (over the PSF) is distributed to a 2D matrix that has the same dimensions as the simulated pixel array. Each element of this matrix represents the number of photons that have arrived at the corresponding pixel during the frame integration period. In step 6.2, shot noise is simulated. It is calculated according to the number of photons counted by the pixel as a Poisson random variable. Notice here that the number of photons that have arrived at each pixel (calculated in 6.1) is different from the number of photons counted as the signal (related to detector quantum efficiency, calculated in 6.2). Next, background noise is added to each pixel in step 6.3. The background is either simulated as an extra component or added from experimental measurements. Lastly, the simulated movie is exported to TIFF stacks in step 6.4 with precision of 16-bit unsigned integer.

The implementations of each step or sub-step are intrinsically independent and interchangeable modules in the simulator. As long as the interface of the module (input and output format) remains the same, they can be replaced, modified, and upgraded independently. Each module can be a function that calculates the output. Alternatively, each module can be an algorithm that simply searches and loads the designated data file based on the input, and then delivers the loaded data file as output. Therefore, the simulator is sustainable and expandable, can be organized and shared in the community, and allows for continuous improvement with collective contributions from the community.

#### *5.2 Detailed explanations of the modules used in this study*

Thanks to the modular design, the simulator can be flexibly upgraded and expanded to account for different optical systems, different sample conditions, etc. As long as the modular interface is maintained, different functionalities can be implemented through the creation of different types of modules. Inputs and outputs for these modules can also be individually manipulated, or handed from one module to the next. Below we explain in more details each existing module included in our package.

Generation of features of interest (module for step 1.1). This module produces a data file with a 2D matrix that represents the locations of all available labeling sites in the simulated sample, representing a chosen morphology. This matrix could represent protein locations with epitopes that could be targeted by immuno-staining, or locations of fluorescence proteins fused to target proteins. The number of rows in the 2D matrix represents the total number of candidate locations (available labeling sites). There are 3 columns in the matrix, each representing the the x-y-z coordinates for each individual candidate emitter location (available labeling sites, or candidate labeling sites). The collection of x-y-z coordinates define the simulated morphology. In this work, we focused on a collection of random filamentous curves, generated by random walk-based Monte Carlo simulations. To generate the curves, we are essentially generating a chain with link points and pre-defined linkage lengths that defines the precision of the simulation. A random initial point is selected as a starting point for the random walk. The direction of the first link of the curve is randomly selected. The algorithm generates a new location which serves as the starting point for the next iteration. The generation of the tilt angle for each step determines the overall appearance of the feature of interest, and four user-adjustable input parameters govern the generation of the tilt angle per each step. A random number is generated to determine the angle tilted for this current step. A provided preference for the angle range of this randomly generated number grants directionality to the simulated curves. Propagating this random walk over all the steps gives us a random curve where the x-y-z position generated at each walk serves as a candidate labeling site. The user can define the step length in order to define the labeling sites density. They can also define the total number of steps in order to define the length of the curve. Generated features of interest consist of the locations of all candidate labeling sites.

'Labeling' available candidate sites (module for step 1.2). This module randomly selects candidate sites (generated in step 1.1) to be occupied by emitters. A user-defined labeling density parameter determines the % occupancy of the sites produced in step 1.1. This module outputs a 2D matrix of occupied sites locations. The total number of rows of this matrix corresponds to the total number of labeled sites. The x-y-z coordinates of these labeled sites are stored in the 3 columns of this matrix.

Updating emitter locations by addition of random errors (module for step 1.3). Coordinates generated in step 1.2. are precisely positioned on the linkage nodes of the chain as the simulated curves. To account for labeling uncertainties, fluorophore linker length, antibody size, etc., emitters locations are displaced from the curve by adding a small random error to the coordinates generated in step 1.2. according to a pre-defined Gaussian distribution with a user-defined width parameter (characterizing the degree of labeling uncertainty). The 2D matrix generated in this step represents the 'ground truth' emitter locations of the labeled sample.

Generating emitters' blinking 'on' / 'off' time periods (module for step 2.1). This module outputs an array of blinking 'on' / 'off' time periods for each individual emitter (to be converted to time trajectories in step 2.2.). Since blinking can be described by a Markov process, these periods are drawn from Poisson distribution functions for 'on' times and 'off' times with $\tau_{on}$ and $\tau_{off}$ as user-defined parameters (the average times the emitter spends in the 'on' and 'off' states respectively). Another user-defined parameter is the photon-emission rate of an emitter in the 'on' state (defined as the 'on' state brightness).

Generating emitters' time-intensity trajectories (module for step 2.2). This module accepts as input the array of blinking 'on' / 'off' time periods (with continues coordinates) generated in step 2.1. and outputs emitters' time-intensity trajectories (with discrete coordinates) by operating on it with a time-integration operator. A user-defined 'frame-duration' parameter is used to bin photons per frame. Each time-intensity trajectory is generated and stored independently for each emitter, forming a library of emitters that can be stored and re-used in other simulations.

Simulation of imaging system PSF (module for step 4). This module yields a 3D matrix in the form of a stack of images that shows the PSF of the imaging system at different focal planes. The computation of the PSF varies with the chosen PSF type as well as the imaging conditions. Note here that this module is independent from the emitter locations and simulated feature of interest. It represents a certain type of imaging system with a certain type of PSF. We use Gibson Lanni's PSF model in this study.

Simulating image formation at the detection plane (module for step 5). This module generates the image formed at the detection plane in the form of cumulated photon-numbers at different spatial locations. The produced data is in the form of a 2D matrix, where the grid size of this matrix defines the precision of this simulation. The image is generated frame-to-frame, and is based on the emitter locations at the given frame, the photon-budget generated by each emitter within the frame, as well as the PSF of the optical system. For a specific emitter, its image at the detection plane is determined by its 3D location coordinates referenced with the defined focal plane. Depending on the distance of this emitter to the focal plane along the optical axis (z-axis), we choose the corresponding PSF image from the pre-computed 3D PSF matrix to be the image of this emitter in the detection plane. The x-y position of this emitter defines the center position of the image of this emitter in the detection plane. The photon signal is then propagated for each photon within the photon-budget of this emitter. The emitter emits a photon, which propagates and arrives at a random position in the detection plane. The arrival location of this photon is randomly generated with a probability distribution described by the image of this emitter (the PSF). To generate the image formed by the simulated sample at the detection plane, we propagate every individual photon generated by every individual emitter of the simulated 3D volume. In general, it describes the spatial distribution of the number of photons that arrives at the detection plane within the time window corresponding to the frame.

Pixilation of the image at the detection plane (module for step 6.1 and 6.2). This module accepts the output from step 5 and projects the data onto the detector grid (camera pixels).

Given the image formed at the detection plane from step 5, we know the spatial distribution of photons that arrive at each camera pixel. For each pixel detector of the camera, we calculate the total number of photons that have arrived at this detector over the given time window. The actual photon counts of this detector are defined by random photoelectron generation process, which is Poisson-distributed. The expectation value (mean) per time frame is the total number of photons that have arrived at this pixel during the frame integration. This allows to independently calculate the shot noise for each pixel (since the noise equals to the mean, and the mean is different for different pixels).

Simulating background noise (module for step 6.3). Aside from the shot noise simulated in module 6.2, background noise is simulated as additive noise in our simulations (through experimental record or simulations). In the case of experimentally measured background noise, a movie of empty frames is acquired by an EMCCD camera (using a blank sample). The number of frames and frame-rate is identical to the simulated movie. The measured movie is then added to the simulated movie, frame-by-frame. Other background signals, such as out-of-focus light or diffusing emitters could also be measured or simulated and added in a similar way.

**Concluding remarks for the simulator**: All modules of the simulator are independent and interchangeable. This flexible design allows us to simulate various morphologies, attribute different photo-physical properties to the emitters, utilize different imaging system (with different point spread functions), and add different background and noise contributions to the signal.

#### *5.3. Specifications of each simulation data.*

There are four sets of simulations used in this study, the first set is the 3 emitter simulations, used in Fig. 3 Fig. S5, S5 and S6. This set of simulation is generated from the same three emitters with blinking statistics depicted in Fig. 3(a) and location depicted with each corresponding figures. The second set is the set of 2 densely labeled lines, used in Fig. S8. For both sets of simulations, the simulated emission wavelength is 520 nm with Gibson Lanni's PSF model, simulated optical system was with 17.78 nm pixel size, which corresponds to 900× magnification if we consider the camera detector pixel size of size 16 μm. This artificially high magnification was selected to avoid binning effect, therefore, isolate the factors that contribute to imperfections in the reconstructed image.

The third set of simulation is a type of simulated feature with randomly distributed curves in 3D. It is used in Fig. 4, Fig. S9, S9, S10 and S11. The dimension of the virtual sample volume is 22.22 μm in the lateral dimensions and 1.8 μm thick. The virtual sample consists of 120 random curves and is populated with fluorophores with different densities and uncertainties as explained in each figure. Due to randomization, the sample naturally possesses dense feature area and sparse feature area, and different FOVs are cropped from the larger sample for comparison of different feature densities as shown in Fig. 4. Variations of the noise-free movies are generated with different labeling densities and labeling uncertainties. Different signal levels were also generated as described in Fig. S11. The full dataset is available on Figshare[9]. Different population of nonspecific labeling fluorophores are generated, and combined with the feature-only movies to simulated different nonspecific labeling conditions as shown in Fig. S10.

The dataset contains 26 simulated movies in total, 14 of them are free of background noise, and 12 of them contains background noise. For the 14 simulated movies free of background noise, we have simulated labeling densities of 20 $\mu m^{-1}$ (with three different labeling uncertainties of 20, 30 and 50 nm), and 50 $\mu m^{-1}$ (with four different labeling uncertainties of 20, 30, 50 and 70 nm). For each of these 7 different combinations of labeling densities and labeling uncertainties, we have simulated two different nonspecific labeling densities of either 2000 or 20,000 nonspecific labels in the field of view. The other 12 simulated movies that contains background noise were derived from the 4 background-free simulated movies with 50 $um^{-1}$ labeling density (with four different labeling uncertainties of 20, 30, 50 and 70 nm). Three signal levels were simulated as outlined in Fig. S11, resulting in 12 simulated movies with 3 different signal levels.

The fourth set of simulation is also random curves. The set of simulated movies were generated from the identical set of simulated labeled samples, but with 100 different focal planes. The simulated labeling density was set to be 50 $\mu m^{-1}$, and labeling uncertainty was set to be 7 nm in order to simulate a sample of very tightly labeled curves in 3D. Photon budget for the emitters were randomly drawn from the range of 1000 to 5000 photons per second at pure 'on'-stage. Background noise was directly added to each frame from an experimentally acquired background noise component as used in the third set of simulations and described in

Fig. S11. The 100 different focal planes result in a datasets that simulated 3D sectioning of the same virtual sample, the result is used in Fig. 5 is available on Figshare[10].

### Appendix 6

In this note we describes the sample preparations for live cell imaging

#### 6.1. Cell Medium

The cell culture medium used in the experiments is composed of 89.29% Dulbecco's Modified Eagle Medium (Gibco 31053-028), 8.93% fetal bovine serum, 0.89% 200 mM L-Glutamine, and 0.89% 100 mM sodium pyruvate.

#### 6.2. Transfection of cells

Wild type HeLa cells were seeded at 10% confluency on a 35 mm glass bottom dish (In Vitro Scientific, D35-20-1.5-N) filled with 2 mL of the cell medium. Cells were allowed to grow for 72 hours up to around 80% confluency in a humidified incubator at 37oC with 5% CO2. For transfection, 3 µL of Lipofectamine 2000 (Invitrogen 11668-019) were mixed with 250 µL of Opti-MEM I Reduced Serum Medium (Gibco 31985-070). 1.2 µg of the DronpaC12-(GGGGS)x3-βActin plasmid (Provided by Prof. Miyawaki's lab) or Skylan-S-(GGGGS)x3-βActin plasmid (Skylan-S protein sequence was provided by Prof. Xu Pingyong's lab, and fused to (GGGGS)x3-β-Actin by ourselves) was also mixed with 250 µL of the Opti-MEM. After 5 minutes of incubation at room temperature, the plasmid mixture was pipetted into the Lipofectamine 2000 mixture and gently mixed by tapping on the Eppendorf tubes. After 20 minutes of incubation at room temperature, 500 µL of this final mixture were used to replace 500 µL of the cell culture medium in the glass bottom dish. Next, cells were returned to the incubator for 5 hours, and old medium was discarded and replaced with fresh 2 mL of cell medium. Cells were left in the incubator for an additional 36 hours.

#### 6.3. Preparation of cells for imaging

To wash away dead cells and cell debris after transfection, the glass bottom dish was washed with 37oC Dulbecco's Phosphate Buffered Saline (DPBS) (Gibco 14190-144). This was done by tilting the dish by approximately 35o, and slowly dripping 10 mL DPBS onto the elevated portion of the dish. While the DPBS run from the elevated part of the dish, a vacuum tip at the bottom removed the DPBS. Cells were kept wet during the entire process. Following, the glass bottom dish was filled with another 2 mL of cell culture medium and placed in the incubator for 3 hours. Medium was then removed from the dish, the dish was re-filled with 37oC 2 mL DPBS, and lightly shaken horizontally. Finally, DPBS was removed, and the dish was filled with another $37^{o}$C 1 mL DPBS (the first addition of DPBS to the cell dish removes most of the cell culture medium which contain trypsin inhibitors).

#### 6.4. Fluorescent imaging of cells

Before imaging, a mixture of 90% DPBS and 10% Trypsin was prepared and heated in a water bath to 37oC. Cells were observed in the open dish (without a lid). Imaging was performed with a wide-field, inverted fluorescence microscope (Nikon Eclipse Ti, Tokyo) with the EGFP filter cube (with 460/60nm band pass excitation filter, 495 nm long pass dichroic and 520/40 nm band pass emission filter) and 485/25 nm excitation (Cyan option, AURA light engine, ©Lumencor, Inc., Beaverton, OR, UCA). Cells with clear actin features were manually selected. When ready to image, 1 mL of the DPBS + Trypsin mixture was dripped on the approximate location of the cell. The field of view was set to 512 × 512 pixels, the EM gain was set to 300, and a 60× objective was used with an extra 1.5× magnifier. The exposure time was set to 0.03 seconds, and 20,000 frame movies were acquired. Light intensity was adjusted to yield brightest pixel values of around ~5000 counts at the start of the data.

#### 6.5. Construction of the gene of Dronpa-C12

A cDNA library was prepared from Acanthastrea sp.; whose tentacles exhibit red coloration. Approximately 150,000 bacterial colonies containing individual cDNA clones were screened for fluorescence. A single clone (R105-8) encoding a cyan fluorescent protein was identified. R105-8 and 22G (the parental clone of Dronpa) were found to be very homologous at amino acid residues between 150 and 176 (89% homology). We made a crossover between the two proteins in the homologous region to generate 22G/R105-8. Then error prone PCR was performed on 22G/R108-8 to create Dronpa-C12. The accession number in the DDBJ/EMBL/GenBank database is LC425653 for Dronpa-C12.

### Appendix 7

This note describes the processing for experimentally acquired data. Images were processed using home-written MATLAB (MathWorks, Natick, MA, USA) codes. Because not every raw data acquired could produce good quality recovery, therefore we will first (explained in 7.1) empirically select from all the raw data for those that are expected to produce good performance for further processing. In 7.2, we explained how we performed data processing for all the identified datasets for further processing of AC2, M6 and so on.

#### *7.1. Empirical selection of raw data for further process.*

First, we inspected $2^{nd}$ order SOFI and $6^{th}$ order SOFI cumulants reconstructed on a randomly selected 400-frame segment of the experimental movie. Next, the two resulting images were manually and empirically inspected to see if the dynamic range of pixel intensities across the full image is acceptable for further processing. Only the usable regions are cropped out for further analysis. There are a few reasons for a region to be discarded that usually appear as abnormally high amplitude of the pixel intensities. The first example is presumably due to broken camera pixels. To be more specific, in the processed cumulant images, a few (usually less than 10) pixels displaying a few orders of magnitude higher than the rest of the pixel values, and the amplitude of these pixels are of orders of magnitude higher than the range of the rest majority of the pixel intensities. In this case, the abnormal pixels are considered un-usable. The second example is a bigger region that (may resemble the shape of a feature of interest) displays abnormally high amplitude of pixel intensities, we would proceed and inspect the acquired movie to see if there are fluorescently labeled features located in the corresponding region that are moving too fast to be resolved at 0.165 Hz (200 fold slower than the original acquisition rate), if confirmed then the region will be discarded. The way to compare whether the dynamics is too fast is initially done by producing two movies from the raw data (acquired at 33 Hz). One movie is produced from the raw data frames where each new frame is the time average of 200 frames from the original raw data frames, and played at an accelerated framerate for convenient inspection (this movie contains 200 fold less frames than the raw data and is expected to display the dynamic process with degraded time resolution). Another movie is produced with the original raw data frames but played at a framerate that is 200 fold faster than the first movie (this movie is expected to display the fast dynamics). Subsequently, these two movies could be played over the same time course side by side for manual inspection. If the dynamics is significantly degraded by motion blur in the first movie with degraded time resolution, then we conclude that the abnormally high pixel intensities in the cumulant is cause by the irregular fluctuations that is greatly affected by the dynamic movement of the feature. In this case, the corresponding portion/region of the raw data will not be included. The third example is the case of photon counts for data acquisition, which often result in smaller or nearly zero fluctuations. This case is identified when a certain region displays abnormally small amplitude of pixel intensities in the cumulant image, with a sharp and abrupt difference with the edges, and very bright pixel intensities in the raw movie. Additional reasons that caused a raw dataset to be discarded could be abnormally high intensity of feature that seemly looks like to be caused by cell debris, or air bubble, with very strong signal but small fluctuations.

#### *7.2. Data process for the selected experimental data.*

All the selected experimental data are divided into 20 continuous segments of 200 frames for further processing. For comparisons, average images, bSOFI analysis, SRRF analysis (with and without deconvSK[1]), $6^{th}$ order SOFI moments (M6) with ldrc (with and without deconvSK[1]) were compared; each producing 20 consecutive processed frames.

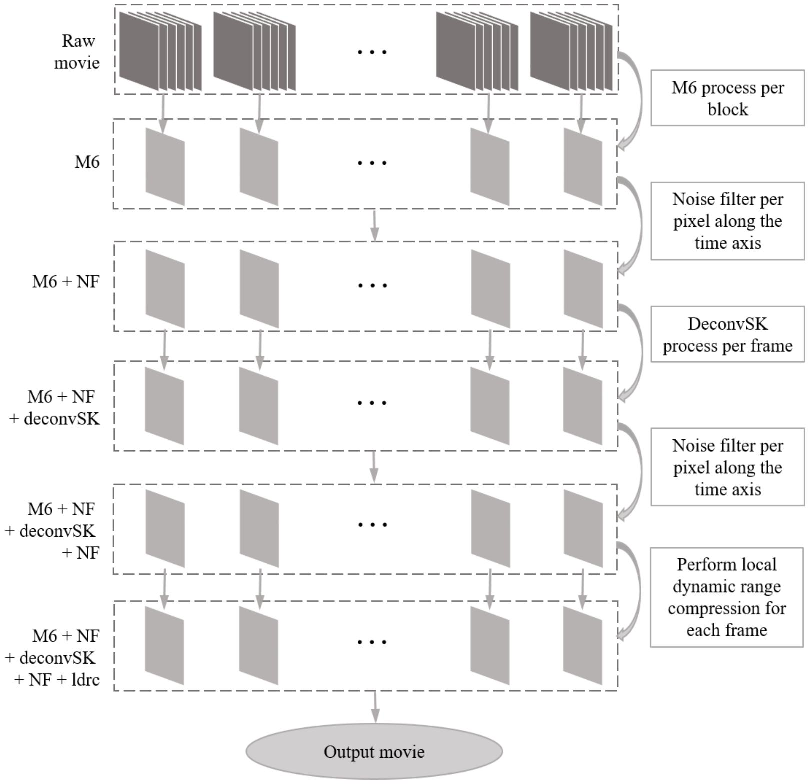

**Fig. S17: Experimental data processing for M6 reconstruction with deconvolution and *ldrc*.** The raw movie (at 30 ms per frame) is divided into segments of 200 frames, where each segment block is processed independently to produce the corresponding M6 reconstruction, and the resultant 20 segments is the new movie at (6 seconds per frame). Each pixel in the field of view is convolved with a Gaussian kernel along the time axis of with width of 2 seconds, filtering out the noise component that contributes faster fluctuations. Note here that the noise filtration for each pixel is independent. DeconSK is then performed on top of the M6 results after noise filtration, producing another movie series with improved resolution. Another layer of noise filtering is added along the time axis to reduce any possible irregular deconvolution artifacts. Local dynamic range compression is then applied for each individual frame with AC2 as the reference image to produce the final output movie.

For the average image, bSOFI and SRRF results, each process is performed on each block individually using direction computation or the open access packages [11, 12]. For the process of M6 images, due to the fact that the resultant images yielded from each individual block is severely degraded by noise, the protocol to filter the noise is explained in Fig. S17. In brief, the raw movie is divided into individual blocks, where each block contains 200 frames. M6 reconstruction is performed per block independently, each yield a single frame for further process. The resultant M6 images yield a 20 frame movie that corresponds to 6 seconds per frame. A noise filter is applied along the time axis for each pixel independently by taking the convolution with a Gaussian kernel with width equivalent to 2 seconds. Deconvolution (DeconvSK [1]) is then applied on each frame of the noise filtered movie to further enhance the spatial resolution. The same noise filter is then applied per pixel along the time axis again to remove any possible irregular deconvolution artifacts. ldrc is then applied on each individual frame to compress the dynamic range of pixel intensities, and produce the final output movie. In principle, the width of the noise filters can be reduced to increase the frequency band in the time domain to allow for better time resolution and customized for different experimental datasets. In our case, we kept every data processing parameter to be the same for convenience.

### Appendix 8

In this note, we provide detailed derivations to achieve (2.17) from equation (2.16) in the main manuscript. We start with equation (2.16) as follows:

$$M1_n(\vec{r};\tau=0)=\sum_{\substack{I_1\cup I_2\cup\cdots\cup I_\sigma\\=\{1,2,...,n\}}}^{\substack{\text{all partitions of}\\\text{set }\{1,2,...,n\}}}\left[\sum_{k=1}^{N}\left(\prod_{p=1}^{\sigma}\epsilon_k^{s_p}\right)\left(\prod_{p=1}^{\sigma}C_{s_p}(\delta b_k(t))\right)U^n(\vec{r}-\vec{r}_m)\right] \tag{SI-8.1}$$

When we look at the first factor in the summation series of (SI-8.1):

$$\prod_{p=1}^{\sigma}\epsilon_k^{s_p}=\epsilon_k^{\sum_{p=1}^{\sigma}s_p}=\epsilon_k^n \tag{SI-8.2}$$

This factor as shown above in equation (SI-8.2) is independent on the partitions. Therefore, in the summation series of (SI-8.1), only the middle factor is partition dependent; the equation can be re-arranged as follows:

$$M1_n(\vec{r};\tau=0)=\sum_{k=1}^{N}\epsilon_k^n U^n(\vec{r}-\vec{r}_m)\sum_{\substack{I_1\cup I_2\cup\cdots\cup I_\sigma\\=\{1,2,...,n\}}}^{\substack{\text{all partitions of}\\\text{set }\{1,2,...,n\}}}\left[\prod_{p=1}^{\sigma}C_{s_p}(\delta b_k(t))\right] \tag{SI-8.3}$$

Furthermore, because we have:

$$M_n\left(\delta b_k(t)\right)=\sum_{\substack{I_1\cup I_2\cup\cdots\cup I_\sigma\\=\{1,2,...,n\}}}^{\substack{\text{all partitions of}\\\text{set }\{1,2,...,n\}}}\left[\prod_{p=1}^{\sigma}C_{s_p}(\delta b_k(t))\right] \tag{SI-8.4}$$

By substituting equation (SI-8.4) into equation (SI-8.3), we get:

$$M1_n(\vec{r};\tau=0)=\sum_{k=1}^{N}\epsilon_k^n U^n(\vec{r}-\vec{r}_m)M_n(\delta b_k(t)) \tag{SI-8.5}$$

Equation (SI-8.5) is then identical with equation (3.5) from the main manuscript. We can see that $M1_n$ is the part representing real emitters at locations $\vec{r}_k$ with virtual brightnessese described by $\epsilon_k^n M_n(\delta b_k(t))$ .

### Supplementary Figure 1

(a)

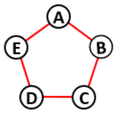

Set of 5 element

(b)

| Number of parts ($q$) | ($f_1$) $f_1 = (-1)^{q-1}(q-1)!$ | ($f_2$) factors for different partitions | |
|---|---|---|---|
| 1 | $(-1)^{1-1}(1-1)! = 1$ | $\langle \delta F_A\, \delta F_B\, \delta F_C\, \delta F_D\, \delta F_E \rangle_t$ | |
| 2 | $(-1)^{2-1}(2-1)! = -1$ | $\langle \delta F_A\, \delta F_D\, \delta F_E \rangle_t \cdot \langle \delta F_B\, \delta F_C \rangle_t$ | $\langle \delta F_A\, \delta F_C\, \delta F_D \rangle_t \cdot \langle \delta F_B\, \delta F_E \rangle_t$ |
| | | $\langle \delta F_C\, \delta F_D\, \delta F_E \rangle_t \cdot \langle \delta F_A\, \delta F_B \rangle_t$ | $\langle \delta F_B\, \delta F_C\, \delta F_E \rangle_t \cdot \langle \delta F_A\, \delta F_D \rangle_t$ |
| | | $\langle \delta F_B\, \delta F_C\, \delta F_D \rangle_t \cdot \langle \delta F_A\, \delta F_E \rangle_t$ | $\langle \delta F_A\, \delta F_B\, \delta F_D \rangle_t \cdot \langle \delta F_C\, \delta F_E \rangle_t$ |
| | | $\langle \delta F_A\, \delta F_B\, \delta F_C \rangle_t \cdot \langle \delta F_D\, \delta F_E \rangle_t$ | $\langle \delta F_A\, \delta F_C\, \delta F_E \rangle_t \cdot \langle \delta F_D\, \delta F_B \rangle_t$ |
| | | $\langle \delta F_A\, \delta F_B\, \delta F_E \rangle_t \cdot \langle \delta F_C\, \delta F_D \rangle_t$ | $\langle \delta F_B\, \delta F_D\, \delta F_E \rangle_t \cdot \langle \delta F_A\, \delta F_C \rangle_t$ |

(c)

$$\begin{aligned} C_5(\delta F_A, \delta F_B, \delta F_C, \delta F_D, \delta F_E) \;=\; & \langle \delta F_A\, \delta F_B\, \delta F_C\, \delta F_D\, \delta F_E \rangle_t \\ & - \langle \delta F_A\, \delta F_D\, \delta F_E \rangle_t \cdot \langle \delta F_B\, \delta F_C \rangle_t - \langle \delta F_A\, \delta F_C\, \delta F_D \rangle_t \cdot \langle \delta F_B\, \delta F_E \rangle_t \\ & - \langle \delta F_C\, \delta F_D\, \delta F_E \rangle_t \cdot \langle \delta F_A\, \delta F_B \rangle_t - \langle \delta F_B\, \delta F_C\, \delta F_E \rangle_t \cdot \langle \delta F_A\, \delta F_D \rangle_t \\ & - \langle \delta F_B\, \delta F_C\, \delta F_D \rangle_t \cdot \langle \delta F_A\, \delta F_E \rangle_t - \langle \delta F_A\, \delta F_B\, \delta F_D \rangle_t \cdot \langle \delta F_C\, \delta F_E \rangle_t \\ & - \langle \delta F_A\, \delta F_B\, \delta F_C \rangle_t \cdot \langle \delta F_D\, \delta F_E \rangle_t - \langle \delta F_A\, \delta F_C\, \delta F_E \rangle_t \cdot \langle \delta F_D\, \delta F_B \rangle_t \\ & - \langle \delta F_A\, \delta F_B\, \delta F_E \rangle_t \cdot \langle \delta F_C\, \delta F_D \rangle_t - \langle \delta F_B\, \delta F_D\, \delta F_E \rangle_t \cdot \langle \delta F_A\, \delta F_C \rangle_t \end{aligned}$$

Fig. S1. A graphical example for how to derive a formula for the 5th order joint-cumulant. (a) Shows a set of five elements (pixels) shown in Fig. 1 in the main text. (b) Shows all possible partitions that contribute to the joint-cumulant. (c) Shows the derived analytical form of this joint-cumulant expressed as a function of individual fluorescence fluctuation trajectories denoted as $\delta F(t)$.

### Supplementary Figure 2

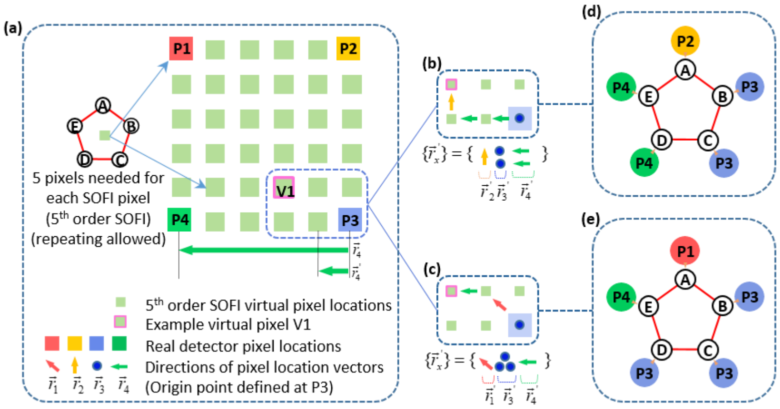

Fig. S2. Example: choice of combination of real pixels for generating 5th order joint-cumulant virtual pixels. (a) Shows a grid of pixels. {P1, P2, P3, P4} indicate real detector pixels. The array of green tiles indicates a finer grid of virtual pixels. $\{\vec{r}_1,\ \vec{r}_2,\ \vec{r}_3,\ \vec{r}_4\}$ is a set of location vectors from P3 (set as the origin) to the real pixels {P1, P2, P3, P4}. Note that in this example, $\vec{r}_3 = 0$. (b) and (c) show two possible paths (and hence real pixel combinations) to reach virtual pixel 'V1' labeled in pink box. (d) and (e) show the corresponding selections of pixel combinations respectively. More details are provided in Supplementary Note 2.

### Supplementary Figure 3

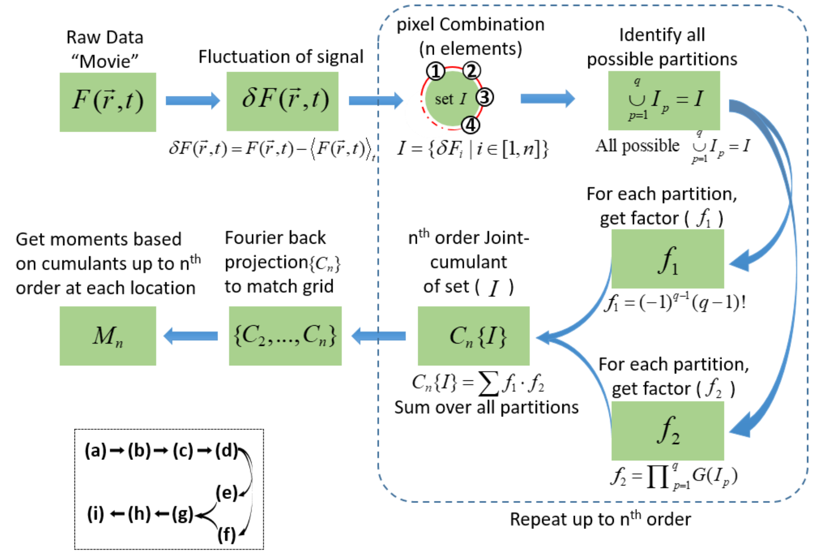

Fig. S3. A flow-chart of the algorithm for joint-moments reconstruction from cumulants. Label for panels (a) – (i) are shown in the bottom left of the figure. The joint-moments reconstruction relies on the joint-cumulants reconstruction. It starts with the (a) raw data (movie) that encodes fluorescence signals as a function of both time and space. (b) Temporal averages are subtracted for each pixel to yield pixel's temporal fluctuation trajectories. (c) to (g) summarize the steps for calculating the cumulants on the (finer) virtual pixels' grid (Fig. S1 and S2), resulting in (h) joint-cumulants reconstruction. For each location (virtual pixel), a full set of cumulants (n = 2 to n = n) is generated. (i) The joint-moments reconstruction is then generated from the full set of cumulants.

### Supplementary Figure 4

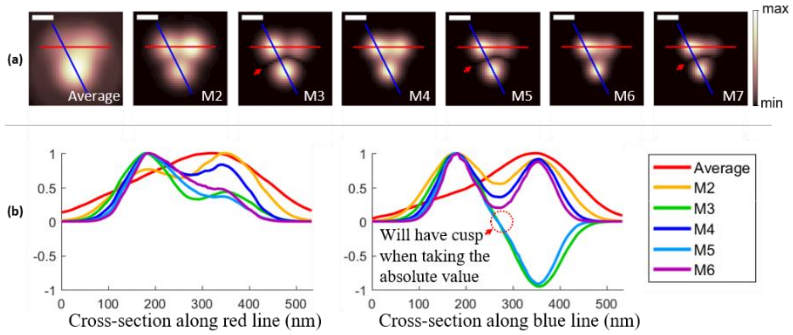

Fig. S4. Cross-section analysis. (a) shows the absolute values of different orders of moments reconstruction as shown in Fig. 3 in the main text. Average image is the average over the entire movie stack, M2~M7 indicates moments of 2$^{nd}$ to 7$^{th}$ order respectively. Each image is displayed with the absolute values at the individual intensity range as indicated in the color bar. Cusps are labeled with red arrows. Two normalized cross-section intensity profiles (as indicated with blue and red lines in different panels in (a)) are plotted in (b). We can see that according to the profile of the average image, the two emitters are not resolved (single peak), but resolved in all orders of moments (two peaks). Additionally, the width of the two peaks in the profiles progressively reduce with the increase of the order of moments, showing the progressive resolution enhancement.

### Supplementary Figure 5

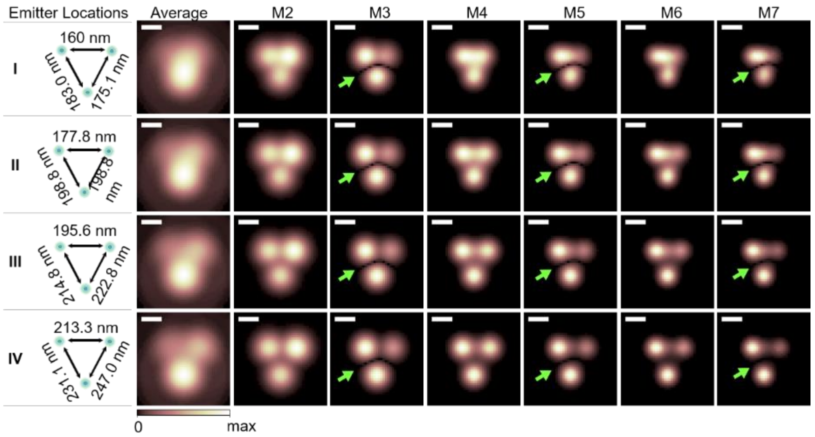

Fig. S5. Moments reconstruction of different orders for 3 emitters positioned on the vertices of an equilateral triangle with different side lengths. The same 3 blinking emitters as used in Fig. 3 are positioned at vertices of different triangles with side lengths I, II, III, IV (indicated in the cartoon, first column). The upper right emitter is successively moved to the right one grid point of 17.78 nm at a time for I, II, III, and IV respectively. The bottom grid point is successively moved downwards one grid point of 17.78 nm at a time for I, II, III, and IV respectively. (Note here that the emitters positions are no longer on the vertices of an equilateral triangle (Note the non-equal distances marked in the cartoons of column 1.) Simulation account for camera pixel size of 16 μm×16 μm, optical magnification of 60×1.5= 90×, corresponding to a grid size of 177.78 nm at the sample plane. However, a 10×10 finer grid (17.78 nm × 17.78 nm) is used in the simulation to simulate sub-pixel resolution requirement. A time series of 4000 frames was simulated with the same blinking parameters and PSF model that were used for the simulations shown in Fig. 3(a) and Fig. 3(b) in the main text. Time averages of the simulated movies are displayed in the second column labeled 'Average'. Moments reconstructions M2 through M7 are shown in columns 3 through 8 respectively. Each image is displayed with its individual intensity range with the universal color map as labeled in the figure. A trend of resolution enhancement as function of order is clearly seen as the reducing size of PSFs. Odd order moments show cusp artifacts (green arrows) while even order moments are free of cusp-artifacts. The larger the triangle's side length is, the better is the reconstruction's fidelity. The displayed images are subject to the individual minimum and maximum pixel values for each panel independently subject to the shown color map in the figure. Scale bars: 160 nm.

### Supplementary Figure 6

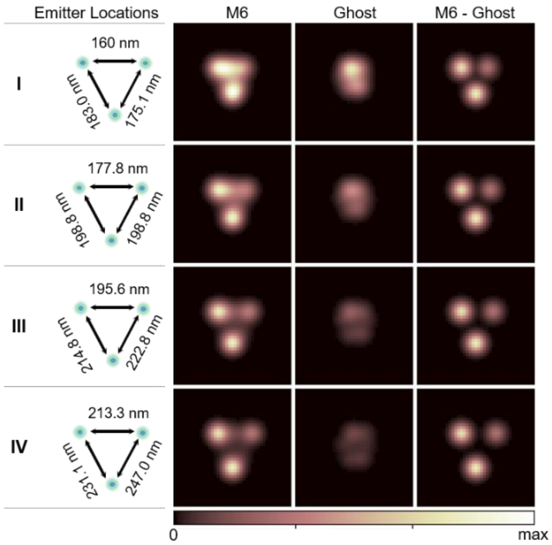

Fig. S6. Decomposing moments reconstruction to virtual emitters located at real emitter locations and non-real locations (‘ghost’ emitters components). Results for the same simulations that were used to construct Fig. S5 are shown again here, but decomposed to ‘ghost’ virtual emitters (located at locations without real emitters), and non-‘ghost’ virtual emitters (located at locations of real emitters), and with their corresponding virtual brightnesses. Emitters locations are shown in the 1st column (same as in Fig. S5). Second column is identical to the ‘M6’ column in Fig. S5, and is calculated from the simulated movie. The third column shows the ‘ghost’ emitters contribution to the images in the second column (M6), and is calculated from the ground truth parameters of the emitters. The fourth column (M6-‘ghost’) is the subtraction of the calculated ‘ghost’ component from the M6 image, showing the non-‘ghost’ virtual emitters contribution to the images in the second column (M6). Note that the ‘ghost’ virtual emitters contribution is greater attenuated with the increase of the distance between the real emitters. The displayed images are subject to the global minimum and maximum pixel values for all panels subject to the shown color map in the figure.

### Supplementary Figure 7

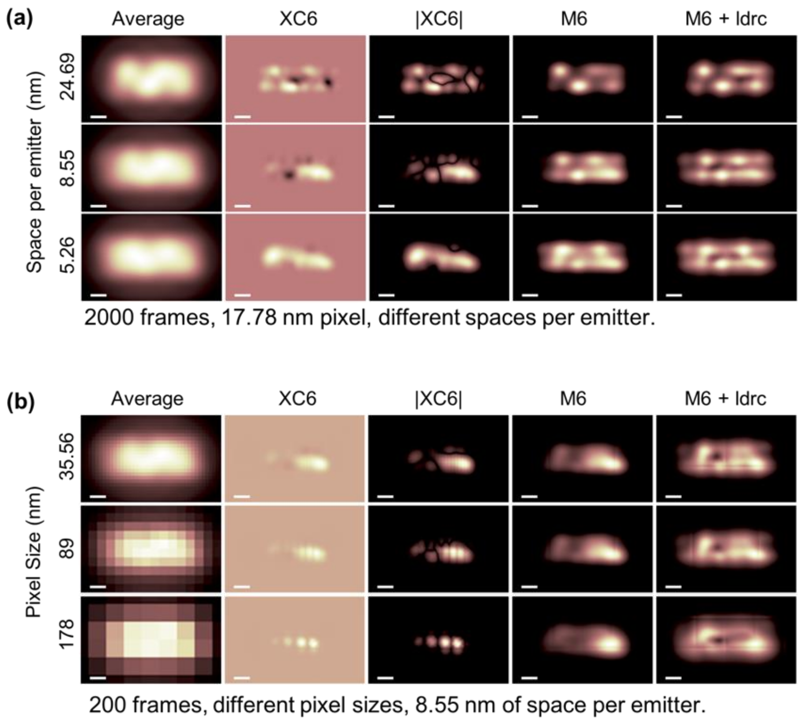

Fig. S7. Comparison between cumulants and moments reconstructions for two near-by filaments (simulated example) and the effect of pixel size. As shown in (a), two filaments (lines) of 889 nm length are placed parallel next to each other 160 nm apart. Emitters are randomly distributed on the lines at 3 different densities of: 24.69 nm/emitter (top row), 8.55 nm/emitter (middle row), and 5.26 nm/emitter (bottom row). Emitters are simulated to emit at 520 nm. Camera pixel size is simulated to be 17.78nm. In (b), the simulated filaments from (a) with 8.55 nm/emitter density (2$^{nd}$ row) is down sampled to yield three different pixel sizes: 35.56 nm, 89 nm and 178 nm. Blinking parameters are randomly assigned. For both (a) and (b), the first column is the time average of the simulated movie. XC6 is the 6$^{th}$ order SOFI (cumulant reconstruction) result with cross-correlations. Note the negative values in this reconstruction. The absolute value of XC6 (|XC6|) is shown in the third column, with noticeable cusp artifacts. M6 (forth column) shows 6$^{th}$ order moment reconstruction results. M6+ldrc (last column) shows results for M6 reconstruction followed by *ldrc* (Appendix 4). The displayed images are subject to the individual minimum and maximum pixel values for each panel independently subject to same color map ('pink' option in MATLAB) as shown in previous figures. Scale bars: 160 nm.

### Supplementary Figure 8

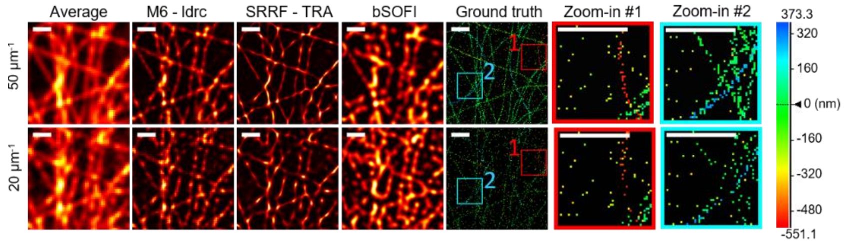

Fig. S8. Comparison between cumulants and moments reconstructions for random filaments with different labeling densities (simulated example)**.** The same field-of-view with two different labeling densities: high density of 50 emitters per μm (top row) and a lower density of 20 emitters per μm (bottom row). The first column is the time average of the simulated movies. M6-ldrc (second column) shows results for M6 reconstruction followed by LDRC. SRRF(TRA) shows SRRF reconstruction with TRA option (third column). bSOFI shows balanced cumulants reconstructed from 2$^{nd}$ to 4$^{th}$ order cumulants (fourth column). Ground truth shows the simulated emitters location in 3D (fifth column) with two zoom-in panels shown as labeled (sixth and seventh column). Emitter height is color-coded according to the color bar on the right. Details of simulation parameters are available in Appendix 5. Scale bars: 640 nm.

### Supplementary Figure 9

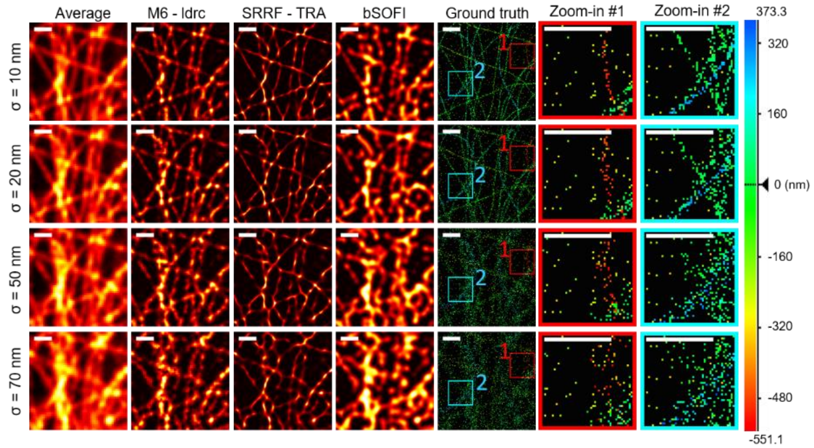

Fig. S9. Comparison between cumulants and moments reconstructions for different filaments labeling uncertainties (or features thicknesses; simulated example). Simulations were performed with different values of labeling uncertainty or feature thickness (σ), of σ = 10, 20, 50, 70 nm (top to bottom rows, respectively). The first column is the time average of the simulated movies. M6-ldrc (second column) shows results for M6 reconstruction followed by LDRC. SRRF(TRA) shows SRRF reconstruction with TRA option (third column). bSOFI shows balanced cumulants reconstructed from $2^{nd}$ to $4^{th}$ order cumulants (fourth column). Ground truth shows the simulated emitters location in 3D (fifth column) with two zoom-in panels shown as labeled (sixth and seventh column). Emitter height is color-coded according to the color bar on the right. Details of simulation parameters are available in Appendix 5. Scale bars: 640 nm.

### Supplementary Figure 10

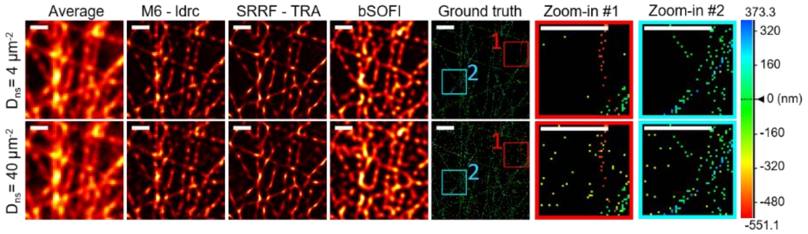

Fig. S10. Comparison between cumulants and moments reconstructions for different nonspecific emitters binding densities (simulated example). Simulations were performed with different densities of emitters non-specifically bound to the cover-slip ($D_{ns}$). Top row shows results for $D_{ns}$ = 4 µm$^{-2}$. Bottom row shows results for $D_{ns}$ = 40 µm$^{-2}$. The first column is the time average of the simulated movies. M6+ldrc (second column) shows results for M6 reconstruction followed by LDRC. SRRF(TRA) shows SRRF reconstruction with TRA option (third column). bSOFI shows balanced cumulants reconstructed from 2$^{nd}$ to 4$^{th}$ order cumulants (fourth column). Ground truth shows the simulated emitters location in 3D (fifth column) with two zoom-in panels shown as labeled (sixth and seventh column). Emitter height is color-coded according to the color bar on the right. M6+ldrc and SRRF(TRA) perform better than bSOFI for this set of simulations. Details of simulation parameters are available in Appendix 5. Scale bars: 640 nm.

### Supplementary Figure 11

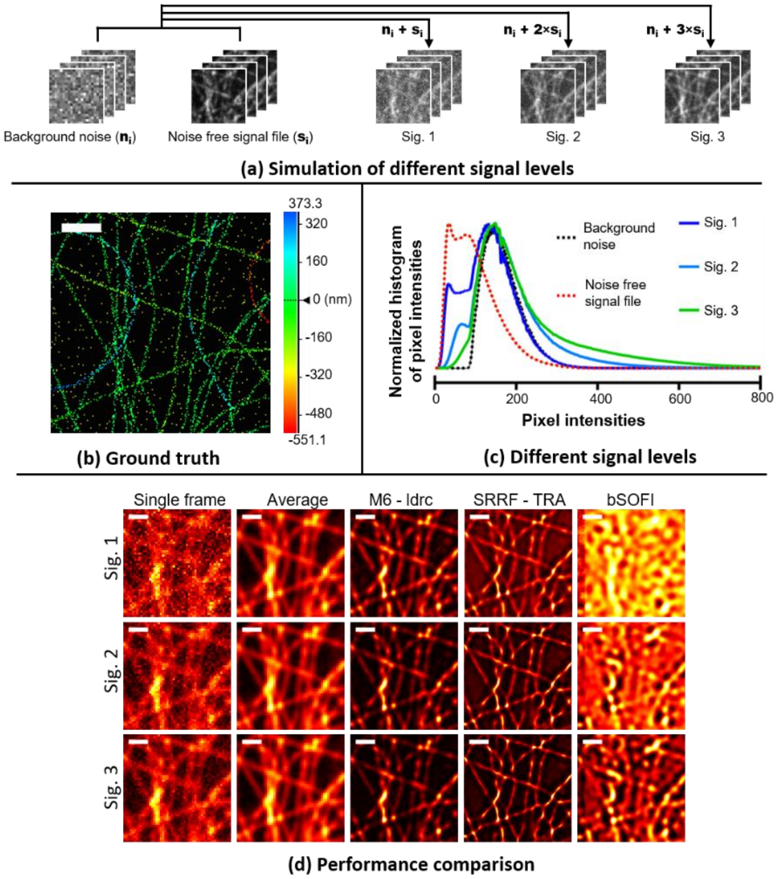

Fig. S11. Comparison between cumulants and moments reconstructions for different signal (to background) levels (simulated example). Simulations were performed with different signal levels. As shown in (a), three different signal levels were simulated with the same background noise. Two separate TIFF movie files were used. One was the *signal file*, a simulated movie free of background noise, denoted as $S_i$ (i stands for the $i^{th}$ frame). The second was the *noise file*, an experimental movie captured by an EMCCD camera a blank sample (without fluorophores), denoted as $n_i$ (i stands for the $i^{th}$ frame). In the constructed noisy movie, each frame is the summation of the corresponding frames from the *noise file* and the *signal file*. 3 different signal levels were generated by multiplying $S_i$ by the factors ×1, ×2, or ×3, resulting in sig×1, sig×2, and sig×3 movies respectively. The ground truth of emitter locations (in 3D, emitter height is color-coded according to the color bar on the right). (b) is therefore identical for all three movies. The normalized histogram of pixel intensities is shown in (c), for *signal file*, *noise file*, and the three different signal level movies. The reconstruction results are shown in (d). The first column shows single frames from the generated movies. The second column is the time average of the simulated movies. M6+ldrc (third column) shows results for M6 reconstruction followed by *ldrc*. SRRF(TRA) shows SRRF reconstruction with TRA option (fourth column). bSOFI shows balanced cumulants reconstructed from $2^{nd}$ to $4^{th}$ order cumulants n (fifth column). M6+ldrc performs better than SRRF(TRA) and bSOFI with respect to S/B. Details of simulation parameters are available in Appendix 5. Scale bars: 640 nm.

### Supplementary Figure 12

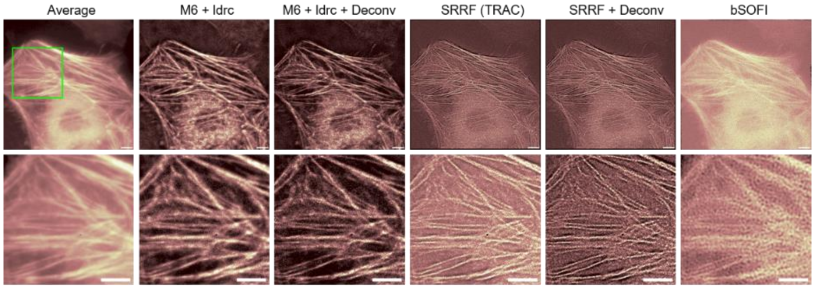

Fig. S12. Comparison of different reconstruction methods for live cell imaging with Skylan-S fused to β-Actin. Reconstruction from live cell 'data-2'. Hela cells were transfected with plasmid that encodes the sequence of Skylan-S fused with β-Actin, and imaged at 30 milliseconds per frame. 200 frames were processed for this figure. Top row is the full field of view, and bottom row is a zoom-in area indicated by the green box. The first column shows the average of 200 frames from the movie 'data-2'. M6+ldrc (second column) shows results for M6 reconstruction followed by *ldrc*. M6+ldrc+Deconv (third column) shows results for M6 reconstruction followed by *ldrc* and deconvSK[1]. SRRF(TRAC) shows SRRF reconstruction with TRAC option (fourth column). SRRF+Deconv shows SRRF reconstruction with TRAC and deconvSK[1] (fifth column). bSOFI shows balanced cumulants reconstructed from 2$^{nd}$ to 4$^{th}$ order cumulants. Scale bars: 8 μm.

### Supplementary Figure 13

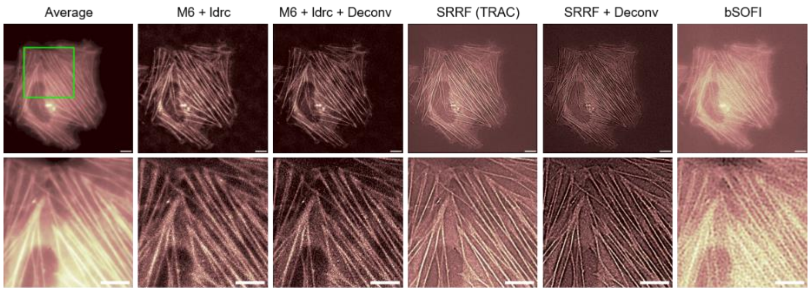

Fig. S13. Comparison of different reconstruction methods for live cell imaging with DronpaC12 fused to β-Actin. Reconstruction from live cell 'data-3'. Hela cells were transfected with plasmid that encodes the sequence of DronpaC12 fused with β-Actin, and imaged at 30 milliseconds per frame. 200 frames were processed for this figure. Top row is the full field of view, and bottom row is a zoom-in area indicated by the green box. The first column shows the average of 200 frames from the movie 'data-2'. M6+ldrc (second column) shows results for M6 reconstruction followed by *ldrc*. M6+ldrc+Deconv (third column) shows results for M6 reconstruction followed by *ldrc* and deconvSK[1]. SRRF(TRAC) shows SRRF reconstruction with TRAC option (fourth column). SRRF+Deconv shows SRRF reconstruction with TRAC and deconvSK[1] (fifth column). bSOFI shows alanced cumulants reconstructed from the 2$^{nd}$, 3$^{rd}$ and 4$^{th}$ order cumulants. (sixth column). Scale bars: 8 μm.

### Supplementary Figure 14

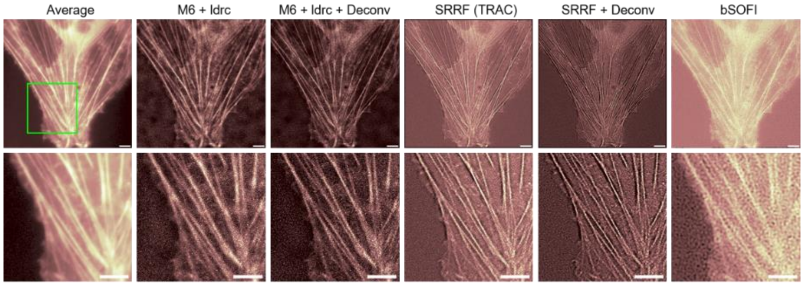

Fig. S14. Comparison of different reconstruction methods for live cell imaging with DronpaC12 fused to β-Actin. Reconstruction from live cell 'data-4'. Hela cells were transfected with plasmid that encodes the sequence of DronpaC12 fused with β-Actin, and imaged at 30 milliseconds per frame. 200 frames were processed for this figure. Top row is the full field of view, and bottom row is a zoom-in area indicated by the green box. The first column shows the average of 200 frames from the movie 'data-2'. M6+ldrc (second column) shows results for M6 reconstruction followed by *ldrc*. M6+ldrc+Deconv (third column) shows results for M6 reconstruction followed by *ldrc* and deconvSK[1]. SRRF(TRA) shows SRRF reconstruction with TRAC option (fourth column). SRRF+Deconv shows SRRF reconstruction (with TRAC option) and deconvSK[1] (fifth column). bSOFI shows the balanced cumulants reconstructed from the $2^{nd}$, $3^{rd}$ and $4^{th}$ order cumulants. (sixth column). Scale bars: 8 μm.

### Supplementary Figure 15

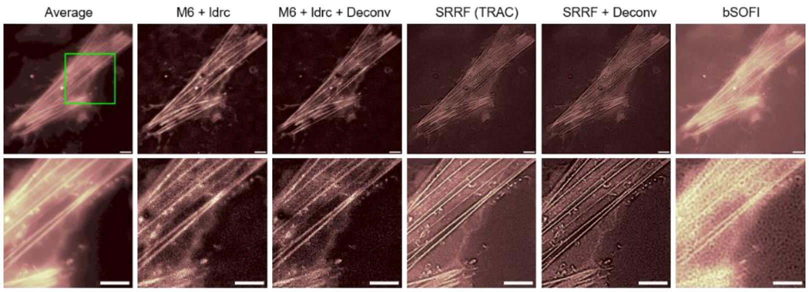

Fig. S15. Comparison of different reconstruction methods for live cell imaging with DronpaC12 fused to β-Actin. Reconstruction from live cell 'data-6'. Hela cells were transfected with plasmid that encodes the sequence of DronpaC12 fused with β-Actin, and imaged at 30 milliseconds per frame. 200 frames were processed for this figure. Top row is the full field of view, and bottom row is a zoom-in area indicated by the green box. The first column shows the average of 200 frames from the movie 'data-2'. M6+ldrc (second column) shows results for M6 reconstruction followed by *ldrc*. M6+ldrc+Deconv (third column) shows results for M6 reconstruction followed by *ldrc* and deconvSK[1]. SRRF(TRA) shows SRRF reconstruction with TRA option (fourth column). SRRF+Deconv shows SRRF reconstruction TRAC and deconvSK[1] (fifth column). bSOFI shows the balanced cumulants reconstructed from 2$^{nd}$ to 4$^{th}$ cumulants (sixth column). Scale bars: 8 μm.

### Supplementary Figure 16

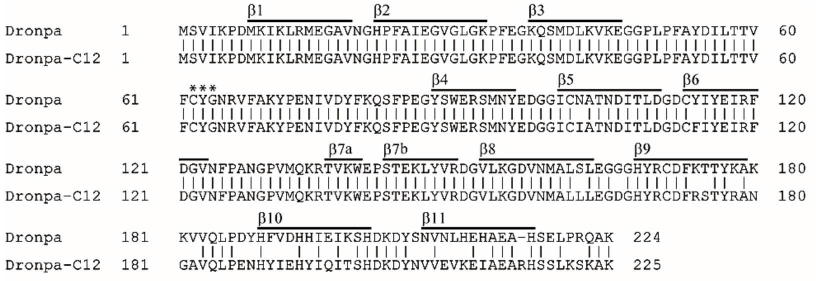

Fig. S16. Amino-acid sequence alignment of Dronpa and Dronpa-C12. The β-sheet-forming regions are represented above the sequences. Residues responsible for chromophore synthesis are indicated by asterisks.
